# Individualised mapping of living human brain mitochondria by MRI reveals signatures of bioenergetic defects

**DOI:** 10.64898/2026.06.09.730804

**Authors:** Michel Thiebaut de Schotten, Cynthia C. Liu, Catherine Kelly, Ke Bo, Tor D. Wager, Eugene V. Mosharov, Martin Picard

## Abstract

Mitochondria support the bioenergetic processes that enable brain function and cognition, but we have lacked a label-free, non-invasive approach to explore how brain mitochondria are linked to ageing, disease, and cognition in humans. A recently introduced MitoBrainMap neuroimaging framework predicts mitochondrial features from magnetic resonance data alone, potentially bridging cellular biology with macroscale brain organization. Here, we tested whether this framework captures meaningful age- and pathology-related mitochondrial variation. Consistent with existing literature, we find that MR-predicted mitochondrial density and tissue respiratory capacity consistently declined with age, whereas mitochondrial respiratory capacity–an index of mitochondrial quality–was relatively preserved across the lifespan. Moreover, the relations among specific mitochondrial features predicted from our algorithm were consistent with their biological organization, supporting preliminary construct validity for MR-predicted mitochondrial features. In patients with rare mitochondrial diseases, predicted maps revealed region-specific alterations in mitochondrial density and respiratory chain components, particularly the expected compensatory upregulation of complex II, but not of other mitochondrial genome-encoded components. Finally, the MR-based mitochondrial features were associated with the energetic stress marker GDF15 measured in blood, as well as with cognitive performance measures, linking the novel predictions of brain mitochondria to systemic stress and behavior. These findings introduce a first-generation, label-free, neuroimaging-based mitochondrial mapping as a non-invasive window into living human brain mitochondria.

---

One and a half billion years ago, an archaeal cell swallowed an aerobic bacterium, establishing a symbiotic relationship^1^. This internalised bacterium replicated and evolved into the mitochondrion, an organelle specialised in converting oxygen and nutrients into membrane potential, used to synthesize cell’s universal energy currency (ATP) and power dozens of other functions^2^. In doing so, mitochondria provided a surplus of energy^3^ and new signaling capabilities^4^. Mitochondrial endosymbiosis thus allowed the host cell to specialise, cooperate with other cells, ultimately leading to the emergence of complex, multicellular organisms with brains specialised to regulate energy metabolism^5^.

Through energy transformation and signaling, mitochondria affect various domains of brain function and cognition, from neurotransmission^6, 7^, neurogenesis ^8^, and social behavior^9, 10^, to cognitive performance^11^. Despite this broad impact, comprehensive mapping of mitochondrial features in humans has long been constrained by methodological limitations, barring recent exciting development^12^, but leaving the brain’s mitochondrial landscape uncharted.

Recently, we made initial progress in overcoming this limitation by biochemically and molecularly profiling a single coronal human brain hemispheric section at a resolution of neuroimaging techniques and developing an MR-based prediction algorithm to predict mitochondrial features across the healthy human brain ^13^. While this approach provided the first spatially resolved atlas of mitochondrial diversity, it remains unclear whether these methods i) can be applied to other MR datasets, and ii) are sufficiently sensitive to capture inter-individual variation in mitochondrial features. If proven to work, this approach could be extended to non-invasive clinical studies to assess mitochondrial biology in individuals with a range of neurodegenerative and neuropsychiatric disorders, and in other contexts.

In humans, mitochondrial diseases are caused by single-point mutations, large-scale deletions, or other defects in mitochondrial DNA. These molecular defects impair oxidative phosphorylation (energy transformation through respiration), leading to multi-system disorders^14^. In its most severe forms, patients develop mitochondrial encephalomyopathy, lactic acidosis, and stroke-like episodes (MELAS), which are often associated with behavioural and cognitive impairments. Mitochondrial diseases also elevate the validated energetic stress marker growth differentiation factor 15 (GDF15)^15, 16^. Genetically defined mitochondrial diseases provide an ideal testbed for validating putative measurements of mitochondrial biology in humans.

Here, we leveraged the Mitochondrial Stress, Brain Imaging, and Epigenetics (MiSBIE) study^17^ to systematically explore novel, label-free MR-based mitochondrial neuroimaging biomarkers in a unique cohort of patients with genetically confirmed rare mitochondrial defects. The neuroimaging portion of the MiSBIE cohort included individuals with single large-scale mitochondrial DNA deletions (n = 10), patients carrying the m.3243 A>G mutation without clinical MELAS (n = 14), and a smaller subgroup with MELAS (n = 2), compared with age- and sex-matched healthy controls (n = 59). We describe the predicted global and regional mitochondrial phenotypes and evaluate their association with age, GDF15, disease severity, and neuropsychological measures.

## Results

As a first step, we replicated the backward linear regression model that predicted post-mortem mitochondrial phenotype^13^ in neuroimaging data from the MiSBIE database^17^. Instead of using a wide sample initially used to develop the model (1,870 university students, ages 18-35^18^), here we retrained a new model on a subset of 10 healthy MiSBIE participants that matched most closely the MitoBrainMap post-mortem brain (54-year-old neurotypical male) in sex and age (males, 36-54 years), and generated diffusion, structural, and fMRI templates (**Supplementary Figure 1**).

**Supplementary Figure 1.**
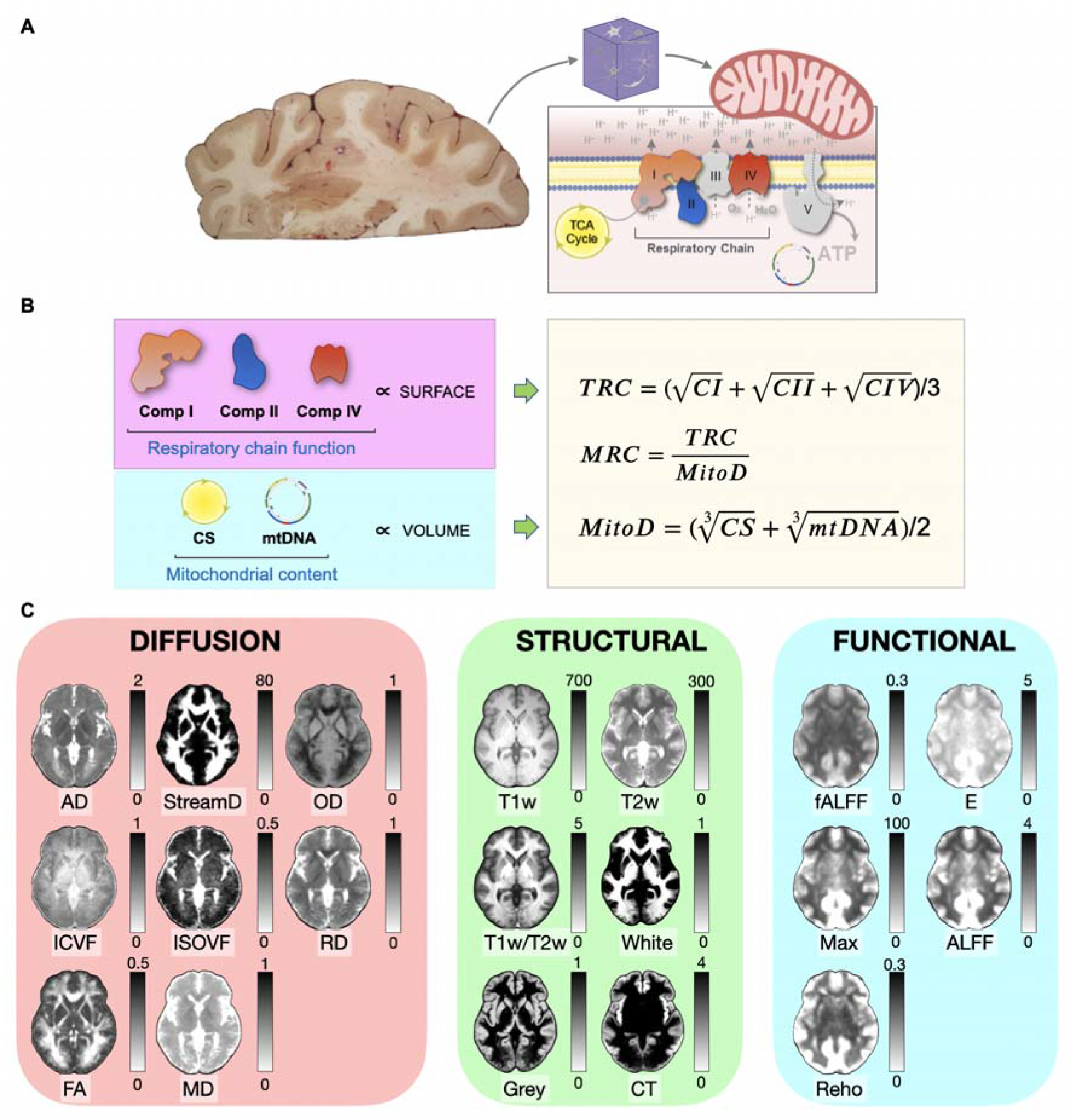
Biological and neuroimaging components underlying in vivo mapping of human brain mitochondrial features. (A) Overview of enzyme activity assays. *Left*, brain slab prior to voxelization.*Right*, schematic of the inner mitochondrial membrane illustrating respiratory chain complexes and ATP production. (B) Conceptual framework linking organelle geometry to mitochondrial function. Activities of respiratory complexes (CI, CII, and CIV) reflect mitochondrial surface-dependent processes, whereas citrate synthase (CS) activity and mtDNA copy number index mitochondrial volume. These measures are combined to derive tissue respiratory capacity (TRC) and mitochondrial density (MitoD). (C) Group-averaged diffusion, structural, and functional MRI templates derived from a subset of 10 healthy participants, approximately matched to the post-mortem brain sample in sex (all males) and age (36-54 years). Diffusion-weighted imaging metrics include axial diffusivity (AD), fractional anisotropy (FA), perpendicular diffusivity (RD), mean diffusivity (MD), streamline density (STreamD), intra-cellular volume fraction (ICVF), isotropic volume fraction (ISOVF), and orientation dispersion index (OD°. Structural MRI metrics include T1-weighted (T1w), T2-weighted (T2w), T1w/T2w ratio, cortical thickness (CT), and probabilistic maps of grey matter (GM) and white matter (WM). Functional %RI metrics include amplitude of low-frequency fluctuations (ALFF), fractional ALFF (fALFF), regional homogeneity (ReHo), maximum BOLD amplitude (Max), and entropy-based measures of signal complexity (E). These multimodal templates served as predictors in the regression models used to estimate voxel-wise mitochondrial features.

Although the new database lacked FLAIR imaging, the linear regression model successfully predicted out-of-sample variance in mitochondrial features, accounting for 20-35% of voxel-level variability (= held-out voxels within the post-mortem section) and 62-79% of regional variability. The predicted features included three electron transport chain enzymes that contribute to energy transformation, complexes I, II, and IV (CI, CII, CIV), mitochondrial density (MitoD), tissue respiration capacity (TRC), and mitochondrial respiratory capacity (MRC), as previously described^13^.

In MitoBrainMap, rather than all mitochondrial features being predicted by a non-descript set of neuroimaging modalities, each mitochondrial feature was predicted by a fairly different set of weights across neuroimaging modalities. These enzyme-specific patterns discovered in the average data thus informed our main hypothesis that MR data contain biological information about specific components of brain mitochondria (total enzymatic capacity, vs. density, vs. mitochondria-specific quality) that could be applied at the individual level.

Applied to MR data templates derived from our 10 new brains, the regression models yielded linear equations that integrate multimodal neuroimaging data to estimate mitochondrial features (see **Supplementary Table 1** for standardized weights, and **Supplementary Figure 2** for comparison with previous predictions^13^. From these, we generated subject-specific mitochondrial measures for the post-mortem slice, and subsequent extension of these predictions across the whole brain based on the same MR-mitochondria model for each individual of our database (**n= 85, Figure 1, Supplementary Figure 3**). To be explicit about the procedure: the regression was fit once, at the template level, by regressing the post-mortem mitochondrial measurements on the group MR templates across the voxels of the post-mortem section. This yielded a single set of standardized weights per mitochondrial feature (**Supplementary Table 1**), obtained after backward elimination selected the most contributing modalities, rather than per-subject coefficients. These fixed weights were then applied to each participant’s coregistered MR maps to generate that individual’s voxel-wise mitochondrial maps.

**Supplementary Table 1.**
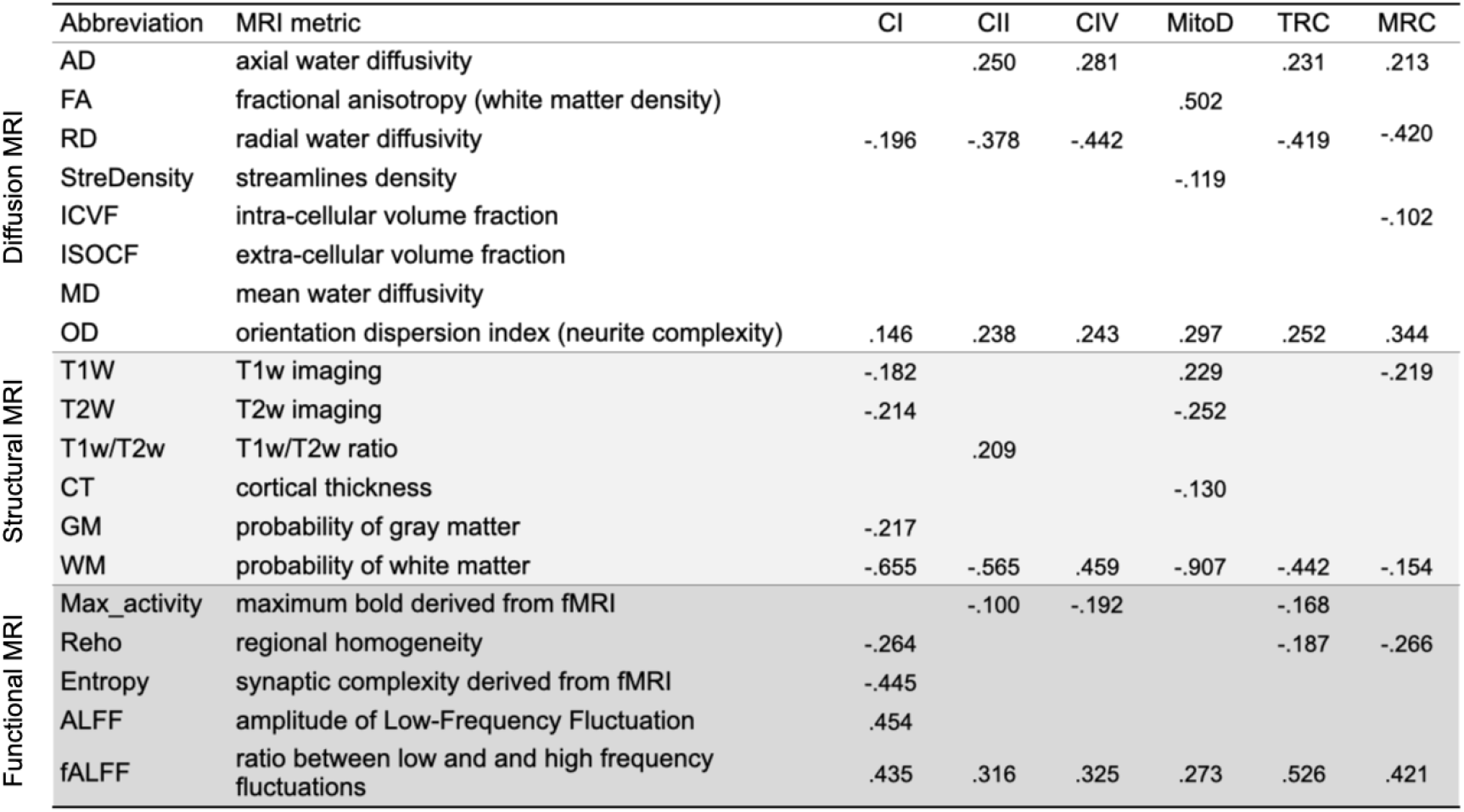
List of neuroimaging metrics and their standardized beta coefficient relationship with the mitochondrial features.

**Supplementary Figure 2.**
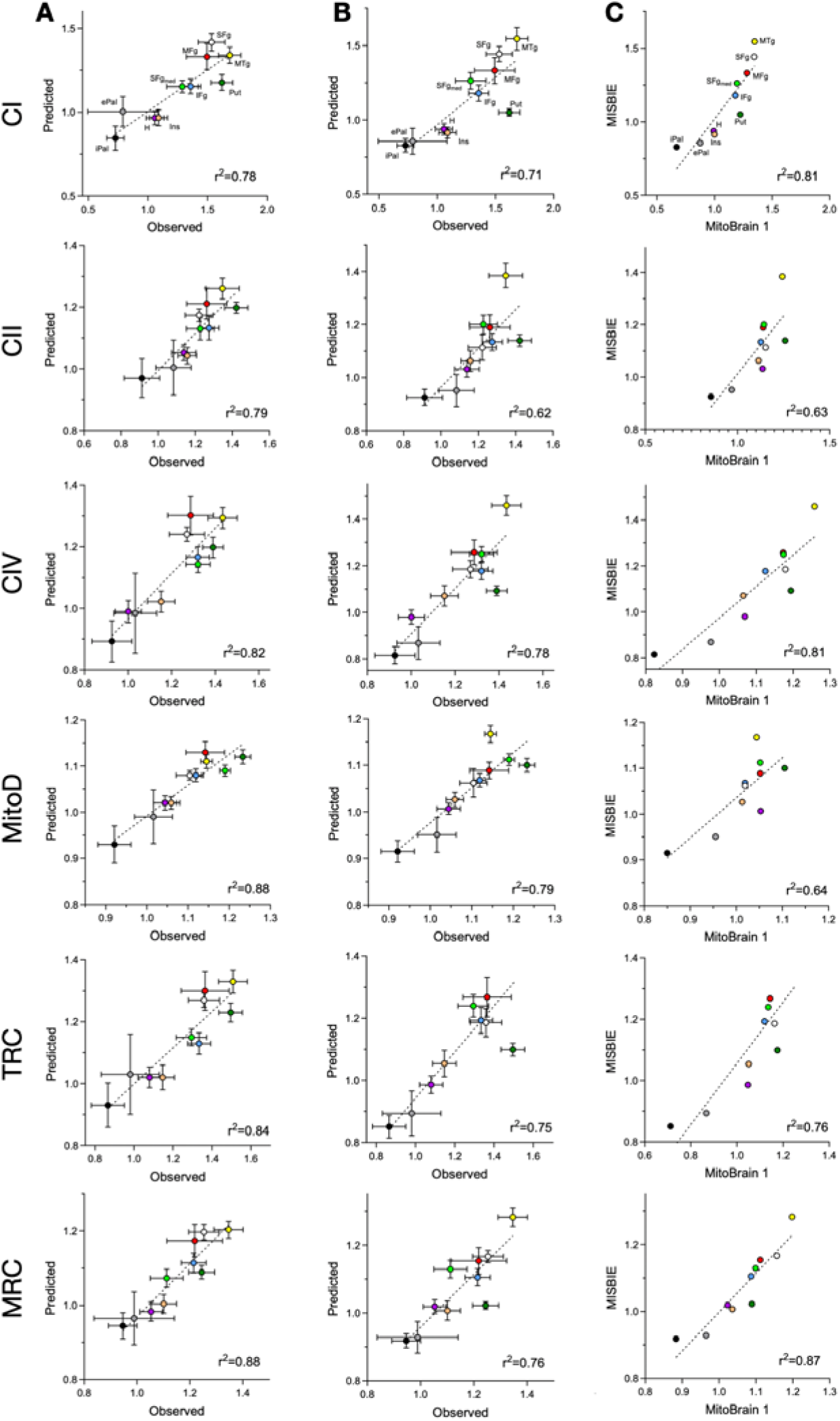
Cross-dataset correspondence of in vivo mitochondrial feature mapping. (A) Regional correlations between observed post-mortem mitochondrial measurements and MRI-predicted mitochondrial features, averaged across cortical and subcortical regions, as reported in Mosharov et al. 2025. Error bars indicate the standard error of the mean across regions. (B) Replication of the same correlations in the independent neuroimaging dataset acquired in the present study. (C) Direct correspondence between regional mitochondrial feature predictions from Mosharov et al. 2025 and those derived from the current dataset, demonstrating cross-study consistency in the predicted mitochondrial maps.

**Figure 1.**
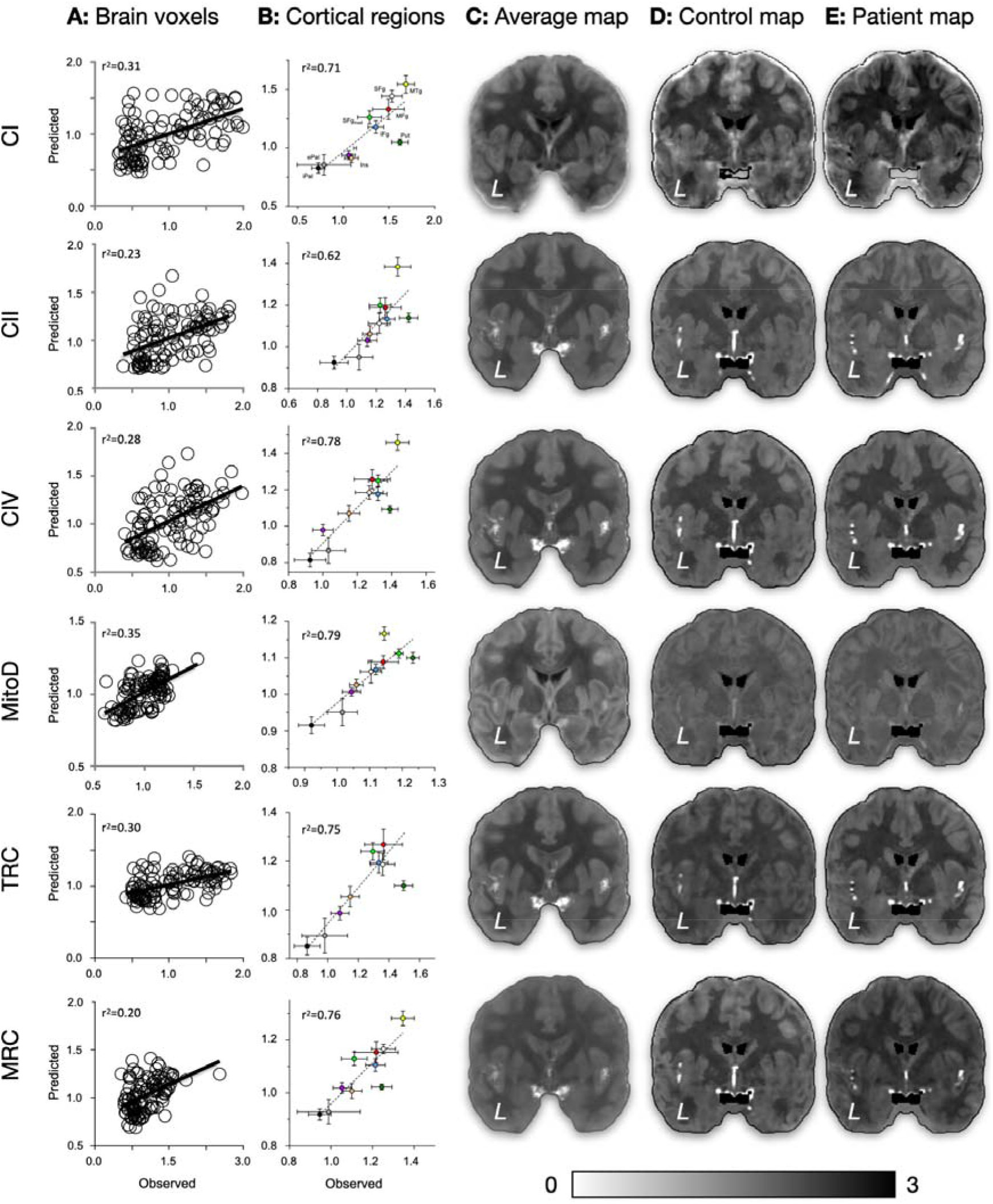
Prediction and mapping of mitochondrial features from neuroimaging data. (A) Voxelwise correlations between out-of-sample observed post-mortem mitochondrial measures and predicted values obtained from the backward linear regression model for Complex I (CI), Complex II (CII), Complex IV (CIV), mitochondrial density (MitoD), tissue respiration capacity (TRC), and mitochondrial respiratory capacity (MRC). Each circle represents a post-mortem voxel, and the coefficient of determination (r^2^) indicates the out-of-sample shared variance between observed and predicted values. (B) Regional correlations between observed and predicted mitochondrial values averaged across cortical and subcortical regions, with error bars indicating standard error of the mean across regions. (C) Representative coronal section of the mitochondrial maps reconstructed from the neuroimaging data of the 10 healthy participants used to train the regression model. (D) Example of predicted mitochondrial maps in an out-of-sample healthy control. (E) Predicted mitochondrial map in a patient carrying a single mitochondrial DNA deletion.

**Supplementary Figure 3.**
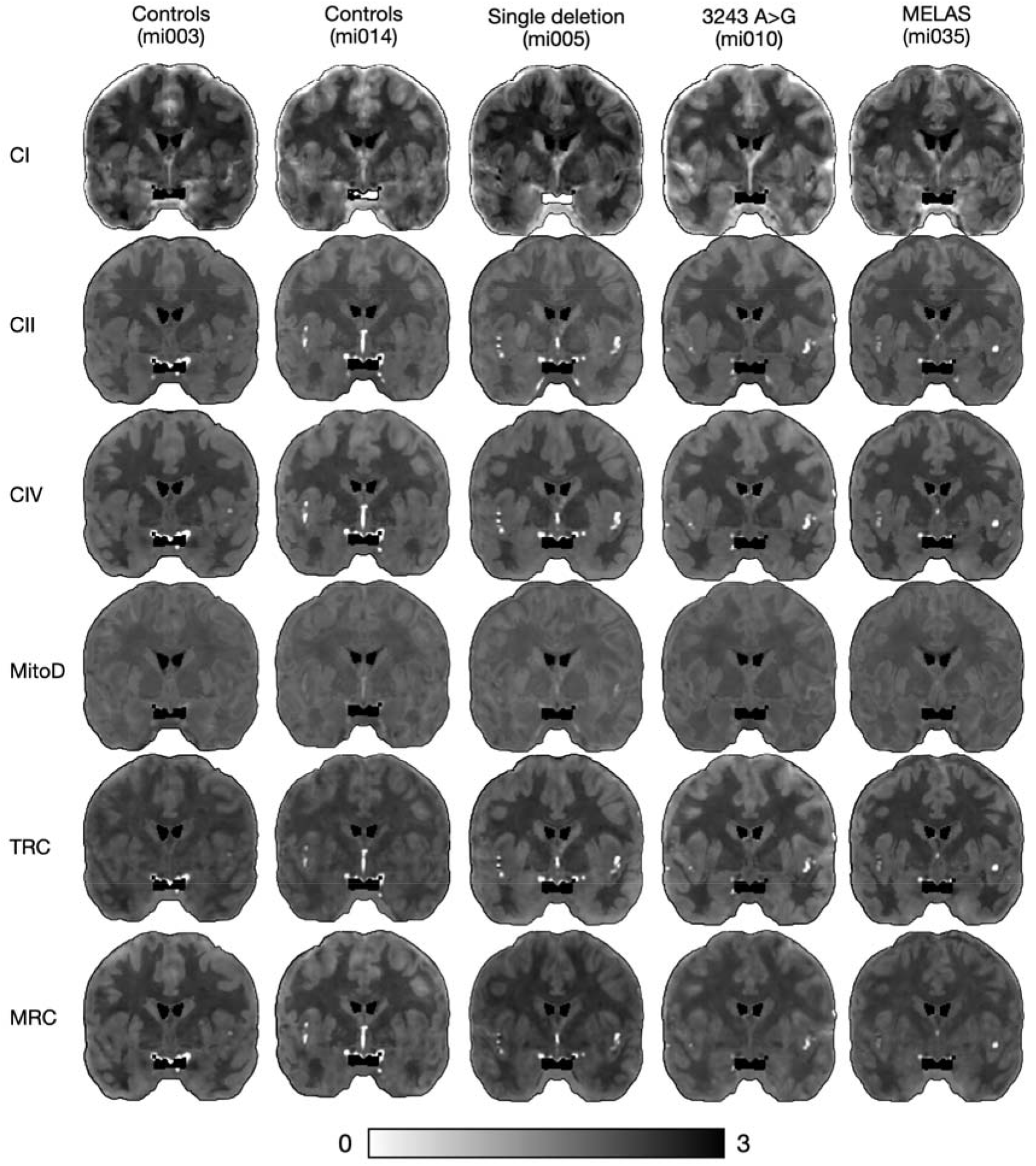
Examples of Controls and patients’ predicted mitochondrial maps

### Age correlations

To evaluate whether individual mitochondrial maps could capture subtle biological changes in mitochondrial features, we examined the well-established age-related decline in mitochondrial density and tissue respiratory capacity ^19-21^; in our whole database (n = 85 subjects), on the slice equivalent to the predicted post-mortem section, while controlling for mitochondrial pathology. We observed robust, negative linear associations between age and both mitochondrial density and tissue respiratory capacity (all P < 0.001), but not the mitochondrial respiratory capacity (**Figure 2A**). These results suggest that between early adulthood and late midlife (20-60 years), mitochondrial content per tissue decreases (MitoD, Pearson r = −0.497 in grey, r = −0.471 in mix and r = −0.417 in white matter, all p < 0.001) reducing tissue respiratory capacity (TRC, Pearson r = −0.526, p < 0.001 in grey matter, r = −0.372, p < 0.01 in mix matter and r = −0.417, p < 0.001 in white matter), whereas the oxidative phosphorylation efficiency of the remaining mitochondria (MRC) is relatively preserved, consistent with age-related changes in skeletal muscle^22^. This finding was further supported by Bayesian analyses showing strong evidence for age effects on CII, CIV, mitochondrial density, and tissue respiratory capacity (all BF_10_> 10), but evidence in favour of the null hypothesis for complex I in white and mixed tissue, and for MRC across all tissue types (BF_10_ ⅓ **Figure 2B**^23^).

**Figure 2.**
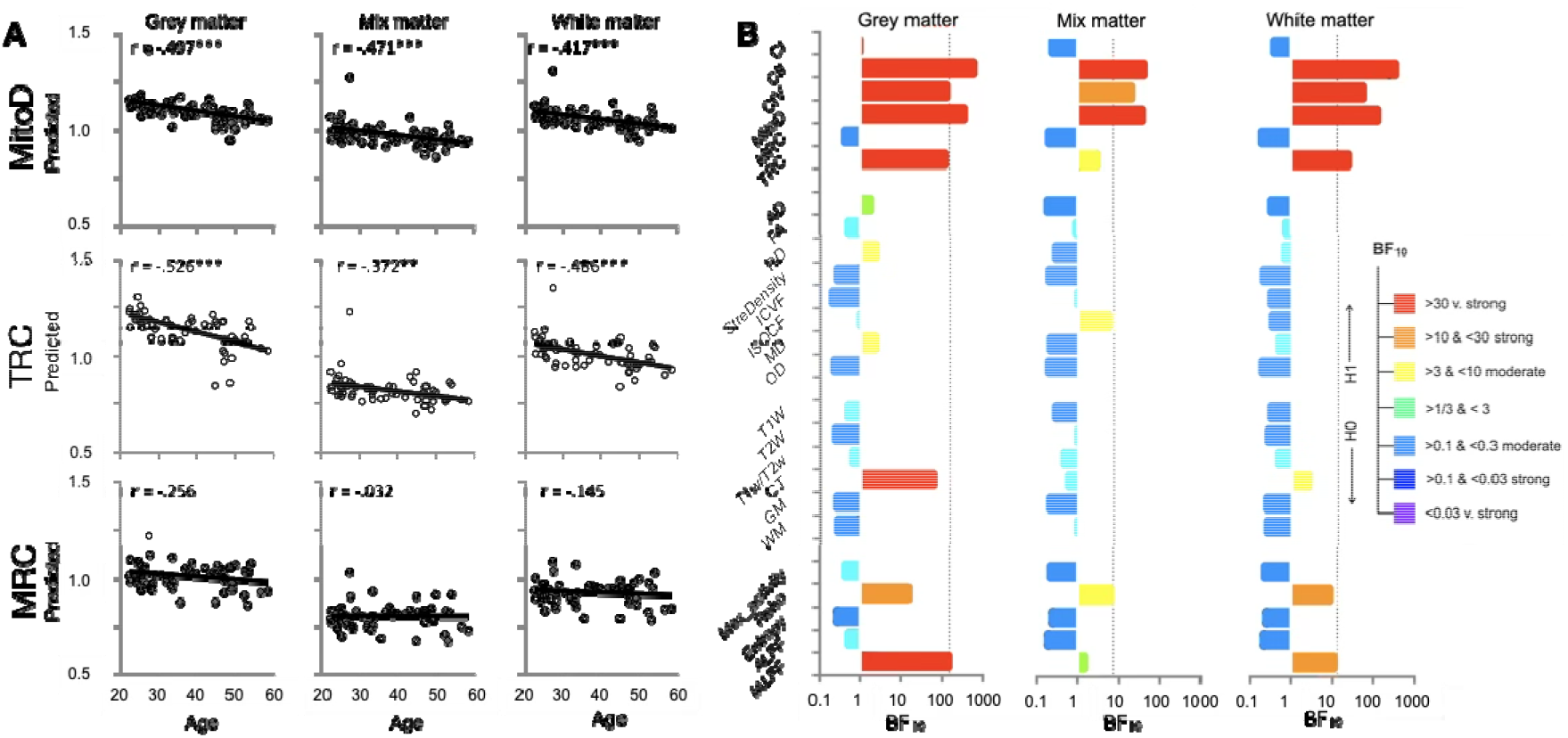
Age effects on predicted mitochondrial features. (A) Predicted mitochondrial features show significant negative correlations with age across participants. (B) Age effects on mitochondrial features exhibit stronger Bayes factors than those observed in any individual neuroimaging modality, indicating that these effects emerge from the multimodal integration rather than a single constituent measure. ** p < 0.01; *** p < 0.001. Bayes Factors (BF_10_) indicate the strength of evidence for the alternative hypothesis (H_1_, significant effect of age) versus the null hypothesis (H_0_, significant absence of effect of age)

Whether the observed age-related effects are specific to mitochondrial features or instead reflect age-related signal changes in one of the MRI modalities contributing to the regression remains unknown. To address this point, we repeated the same analysis separately for each MRI modality and compared the resulting Bayesian values (B_10_) with those obtained for the computed mitochondrial features. Results indicated that, across modalities, the age effects derived from the mitochondrial maps consistently yielded higher Bayes factors than those of any individual modality, suggesting that the integrated mitochondrial phenotype captures biologically relevant variance beyond that explained by each neuroimaging modality alone (**Figure 2B**). Here the grey, white, and mixed values are averages across the entire corresponding tissue compartment of the predicted slice, and the competing predictors were precisely the constituent MR metrics entering the mitochondrial models, evaluated on the same voxels. As an additional control, a simple unweighted average of all input MR maps did not correlate with age in any tissue compartment (Pearson r = −0.036, BF_10_ = 0.143 in grey matter, r = 0.010, BF_10_ = 0.136 in mix matter and r = −0.103, BF_10_ = 0.209 in white matter), confirming that the age effect emerges specifically from the mitochondrial weighting rather than from the aggregation of the underlying modalities.

### Disease contrasts

Next, we examined whether patients with genetically confirmed mitochondrial disease (single deletion, MELAS, and m.3243 A>G mutation without clinical MELAS) exhibit quantitative shifts in specific mitochondrial features at the tissue level on the slice equivalent to the predicted post-mortem section.

We first explored these differences by means of standard multivariate analysis of covariance (MANCOVA, with age as covariate, **Figure 3A**). This analysis revealed significant groups differences for complex II (CII; grey matter: F_(3, 80)_ = 4.479, p = .006; mixed matter: F_(3, 80)_ = 4.730, p = .004; white matter: F_(3, 80)_ = 4.818, p = .004), mitochondrial density (MitoD; grey matter: F_(3, 80)_ = 3.581, p = .017; mixed matter: F_(3, 80)_ = 4.513, p = .006; white matter: F_(3, 80)_ = 4.465, p = .006) and tissue respiratory capacity (TRC; grey matter: F_(3, 80)_ = 2.853, p < .042), but not for Complex I or mitochondrial respiratory capacity (BF_10_< 1/3). The group effects remained significant after exclusion of the small (n=2) MELAS subgroup (see **Supplementary Table 2**).

**Figure 3.**
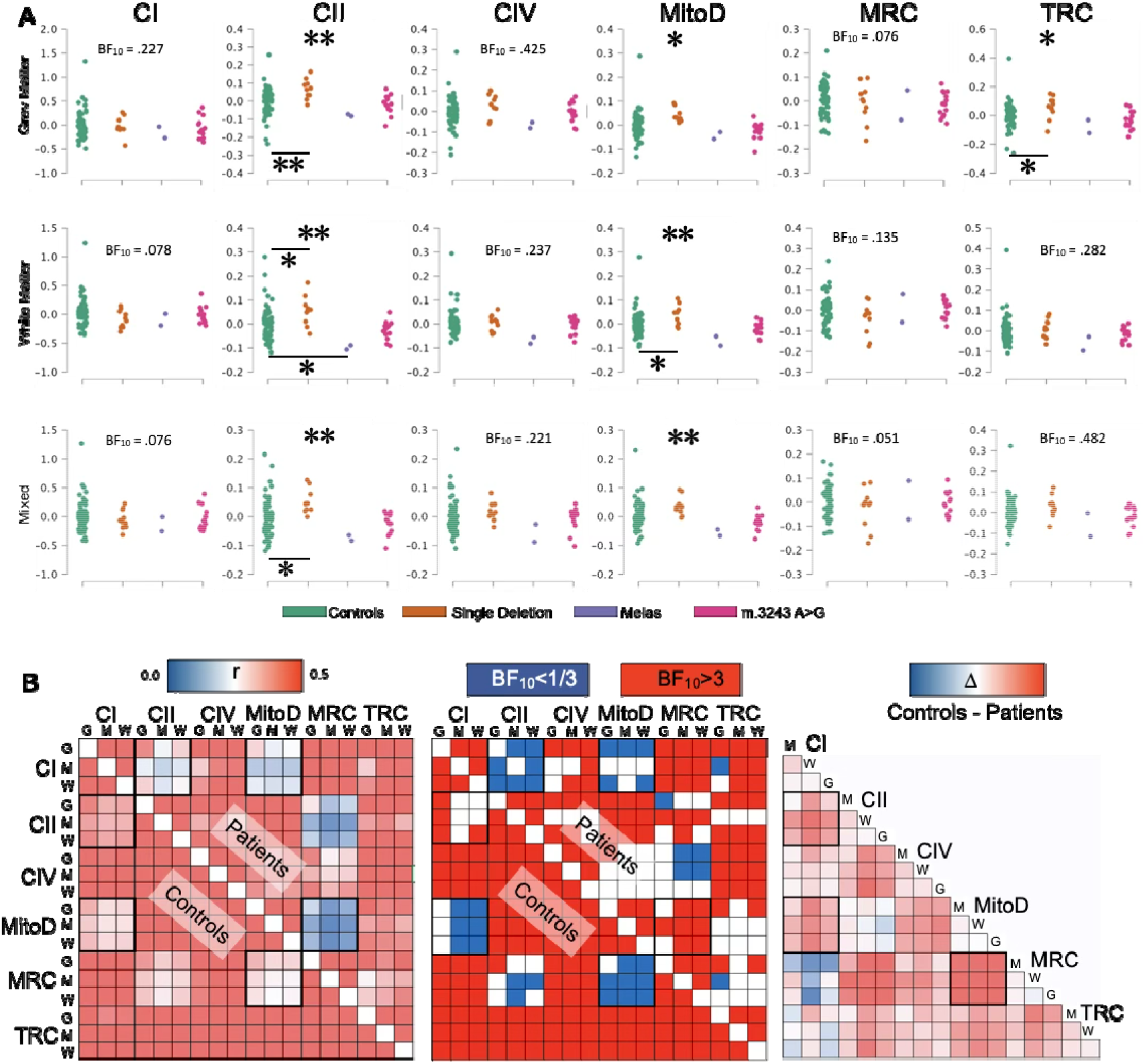
Group effects on predicted mitochondrial features. (**A**) Predicted mitochondrial features differ between controls and patients with genetically confirmed mitochondrial disease, even after age correction. CI: complex I; CII: complex II; CIV: complex IV; MitoD: mitochondrial density; MRC: mitochondrial respiratory capacity; TRC: tissue respiratory capacity; * p < 0.05; ** p < 0.01; *** p < 0.001. Bayes Factors (BF_10_) indicate the strength of evidence for the alternative hypothesis (H_1_, significant effect of age or mitochondrial disease) versus the null hypothesis (H_0_, significant absence of effect of age or mitochondrial disease). (**B**) Coordinated bioenergetic regulation (i.e., homeostasis) across mitochondrial features and groups. *Left*, correlation matrices of predicted mitochondrial features reveal a structured pattern of inter-feature coupling in controls (lower triangle) and patients with genetically confirmed mitochondrial disease (upper triangle), consistent with coordinated, homeostatic regulation across respiratory complexes, mitochondrial density, and tissue respiratory capacity. Each row or column represents Pearson’s r coefficients for the correlation of between indicated mitochondrial modalities in grey (G), white (W) or mixed (M) matter. *Middle*, Bayesian analysis of correlation strength identifying pairs of preserved coordinations (BF_10_ > 3 red) or altered coordination (BF_10_ < 1/3, blue). *Right*, Difference matrices (controls–patients), highlighting selective disruption of this homeostatic coordination beyond shifts in magnitude.

An exploratory post hoc analysis was conducted to further characterize group effects. To limit the number of comparisons, pairwise tests were restricted to contrasts against the control group when the main effect of the group was significant (p-values are Bonferroni-corrected; n = 48^24^). This corresponds to a protected post-hoc procedure, in which pairwise comparisons are conducted only when omnibus group effect is significant; gating the contrasts on a significant omnibus test controls the family-wise error rate, and the comparisons were additionally Bonferroni-corrected. Restricting contrasts to the control reference group is hypothetically driven and reduces, rather than inflates, the number of tested comparisons. As shown in **Figure 3A**, patients with a single large-scale mtDNA deletion exhibited higher CII values in grey (p = 0.003), white (p = 0.013) and mixed matter (p = 0.006), as well as higher mitochondrial density in grey (p = 0.019), white (p = 0.020) and mixed matter (p = 0.014), and higher tissue respiratory capacity in the grey matter (p = 0.046). The selective upregulation of complex II is a biochemical feature of mitochondrial diseases caused by the fact that complex II is uniquely encoded in the nuclear genome, unlike complexes I and IV that are affected by mtDNA defects^25-29^. Notably, the tissue-level activities of the mtDNA-dependent complexes I and IV did not decrease in patients relative to controls. This is expected rather than contradictory: the predicted features index respiratory-chain activity per unit of tissue, so the impaired per-mitochondrion function of complexes I and IV is offset by the compensatory increase in mitochondrial content, leaving their tissue-level activity broadly unchanged. The disease signature detectable here is therefore a redistribution among components corresponding to an elevated nuclear-encoded complex II together with increased density, rather than an absolute fall in complex I and IV.

In contrast, given the small sample size, patients with MELAS appeared to have lower CII in the white matter and a trend toward reduced mitochondrial density, consistent with the previous observation, in immune cells, that individuals who fail to upregulate mitochondrial content develop more severe disease, whereas those who increase mitochondrial density are protected^30^.

**Supplementary Table 2.** Group effects (F statistics from MANCOVA), corrected for age, for each mitochondrial feature across the grey, white, and mixed matter, excluding the MELAS group, with corresponding p values.

|  | Mito Feature | F | p value |
| --- | --- | --- | --- |
| Grey Matter | CI | 0.65 | 0.58 |
|  | CII | 4.48 | 0.01 |
|  | CIV | 1.81 | 0.15 |
|  | MitoD | 3.58 | 0.02 |
|  | MRC | 0.40 | 0.76 |
|  | TRC | 2.85 | 0.04 |
| White Matter | CI | 0.81 | 0.49 |
|  | CII | 4.82 | 0.00 |
|  | CIV | 1.24 | 0.30 |
|  | MitoD | 4.47 | 0.01 |
|  | MRC | 1.61 | 0.19 |
|  | TRC | 1.21 | 0.31 |
| Mixed | CI | 0.47 | 0.70 |
|  | CII | 4.73 | 0.00 |
|  | CIV | 1.09 | 0.36 |
|  | MitoD | 4.51 | 0.01 |
|  | MRC | 0.59 | 0.62 |
|  | TRC | 1.85 | 0.15 |

### Coupling of mitochondrial features

Mitochondrial features are expected to be statistically coupled because they reflect coordinated bioenergetic regulation and organization within the organelle^31, 32^. Consistent with this biological reality, correlation analyses among MR-predicted features revealed a structured pattern of associations across mitochondrial complexes, density, and respiratory capacity in controls. Specifically, CII and MitoD, which are encoded exclusively by the nuclear genes, as well as CI and CIV, which require mitochondrial transcription, were all positively correlated (**Figure 3B left panel**).

If all mitochondrial features were randomly predicted by MR-based parameters, the correlation structure among all mitochondrial features in healthy controls would be expected to be either random (some positive, some negative, if features were noisy), or uniformly positive (if all mitochondrial features were spuriously related). Our results showed no negative correlations and displayed a lack of correlations between MRC and nuclear-encoded markers of mitochondrial content (complex II and MitoD). This signature was also consistent among three distinct brain tissue types (grey, mixed, white), We note that, because the underlying MR maps are themselves spatially structured, some organised correlation among predicted features would be expected even under arbitrary weights; the mere presence of structure is therefore not, on its own, strong evidence. What is informative is, however, the specificity of the pattern, in particular the selective decoupling of MRC from the nuclear-encoded markers of content, and more decisively, the biologically predicted reconfiguration of this structure in patients (next paragraph), which arbitrary weightings would not be expected to reproduce. With this caveat, the alignment of the control correlation structure with known mitochondrial biology is consistent with these features carrying genuine biological information.

In contrast, in mitochondrial disease participants where mtDNA is affected, mitochondrial respiratory capacity was not correlated with complex II nor with MitoD. This signature suggests that individuals with a range of mitochondrial quality (MRC) achieve a relatively stable tissue respiratory capacity (the key parameter to subserve brain bioenergetics) by altering brain mitochondrial content–through compensatory mitochondrial upregulation.

As an additional validity test, we leveraged the well-known process of compensatory upregulation of nuclear-encoded complex II and mtDNA in mitochondrial diseases^30^. When the mtDNA is defective, it adversely affects mtDNA encoded complexes I and IV that combine into the MRC, decreasing these components in human tissues ^25-29^. Based on these biologically-driven relationships, we hypothesized that this coupling would be altered in patients with genetic mitochondrial defects.

As predicted, the Bayesian model provided evidence for the absence of association for most CI–CII and CIV–MRC feature pairs, alongside evidence for a negative relationship between mitochondrial density and respiratory capacity (**Figure 3B midle pannel**). Mitochondrial disease patients with lower MRC (more severe energetic deficit) exhibited the highest mitochondrial content. This response is normally seen in skeletal muscle of affected patients^33^ and in individual cells with mtDNA mutations and deletions^34^. A difference matrix (**Figure 3B right pannel**) confirmed the disruption of these interdependencies (upregulation of CII and mitochondrial mass in CI-deficient tissue) and the reweighting of specific feature relationships (MitoD-MRC) specifically in individuals with mitochondrial diseases.

Together, these findings indicate that genetically-driven mitochondrial respiratory chain defects are associated with a reconfiguration of inter-feature coupling, reflecting the compensatory stabilization of tissue respiratory capacity through the upregulation of nuclear-encoded mitochondrial biogenesis. To our surprise, these data show that this enzyme-specific biochemical pattern is reflected by MRI-predicted features.

### Brain-wide analyses

To determine whether age-related and genetic mitochondrial alterations exhibit brain-area-specific profiles, we applied the same statistical models voxel-wise across the whole brain, with family-wise error (FWE) correction for multiple comparisons. Age effects were not uniformly distributed, predominantly affecting MRC and TRC in subcortical regions (including the pallidum, putamen, hippocampus, and amygdala) as well as cortical territories such as the orbitofrontal cortex, the hand-knob region, and the visual word-form area. In contrast, major white matter pathways, including the optic radiations and the corpus callosum, appeared less affected (**Figure 4A**).

**Figure 4.**
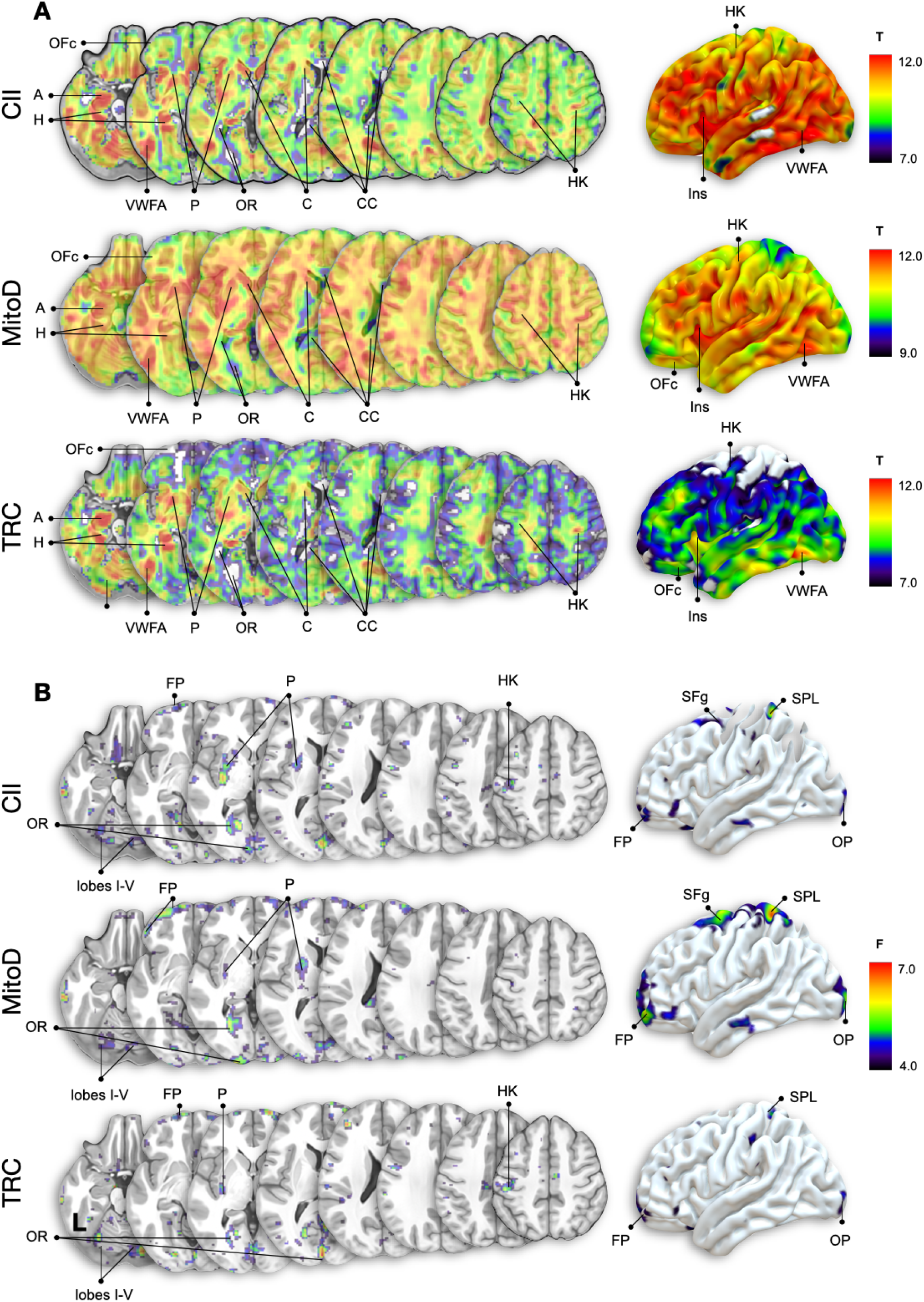
Whole brain age- and disease-related differences in the predicted mitochondrial features. (A) Voxel-wise maps showing the effect of age on predicted complex II (CII) mitochondrial density (MitoD) and tissue respiratory capacity (TRC) across all participants. Warmer colours indicate stronger age-related decreases with peaks in the orbitofrontal cortex (OFC), insula (Ins), caudate (C), putamen (P), visual word for area (VWFA), hand knob area (HK), with a relative preservation of the whole corpus callosum and the optic radiations. (B) Whole-brain group comparison (F values, MANCOVA, age regressed out) for CII, MitoD, and TRC between mitochondrial disease patients and neurotypical brains. Although quite modest in their spatial extent, these maps display significant clusters in the frontal pole (FP), superior frontal gyrus (SFg), left-hand knob area (HK, controlling the right hand), orbitofrontal cortex (OFc), putamen (P), superior parietal lobule (SPL), occipital pole (OP), and optic radiations (OR), as well as in the cerebellum (lobes I-V). Warmer colours indicate stronger disease-related increases in mitochondrial parameters. As significant age and group-related effects were only identified for CII, MitoD, and TRC, voxel-wise analyses were restricted to these mitochondrial indices to reduce the number of comparisons, avoid incidental findings, and focus on the most biologically relevant features. Functional labeling of task-relevant cortical regions was supported by independent meta-analytics of activation maps (**Supplementary Figure 4**)

Mitochondrial genetic defects appeared to target specific regions, with significant group differences in complex II (CII), mitochondrial density (MitoD), and tissue respiratory capacity (TRC) in the orbitofrontal cortex, hand-knob region, posterior putamen, and optic radiations (**Figure 4B**), but not in other regions.

**Supplementary Figure 4.**
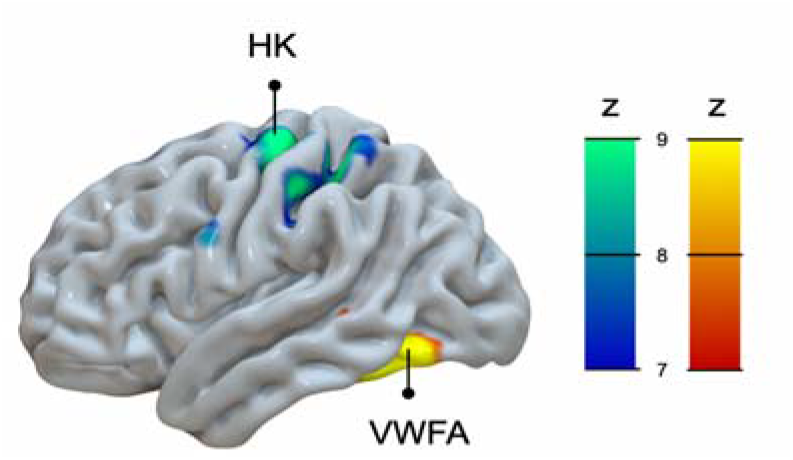
Typical functional activations of the hand-knob and visual word form area. Meta-analytic activation maps illustrate typical cortical responses associated with hand movement tasks, highlighting the hand knob region (in cyan, HK) and reading tasks, activating the visual word form area (in yellow, VWFA). Maps were derived from the Neurosynth meta-analytic database (https://neurosynth.org) and displayed as Z-score maps summarizing consistent task-related activations across studies^35^.

### Brain-biomarker associations

Patients with mitochondrial genetic defects report elevated fatigue, higher perceived stress levels, and reduced stress tolerance^16, 36, 37^. We therefore complemented our neuroimaging markers of mitochondrial biology with a systemic biomarker of mitochondrial energetic stress, plasma GDF15^15, 16^.

In the MiSBIE study patients, plasma GDF15 levels were sampled at nine time points, including a fasting baseline collected before breakfast, a pre-stress measure obtained 5 minutes before the stress induction (a brief oral presentation delivered to an evaluator), and subsequent samples acquired 5, 10, 20, 30, 60, 90, and 120 minutes after the stress exposure. As summarised in **Figure 5A** and **Supplementary Table 3**, GDF15 levels showed moderate evidence for an association with higher mitochondrial density in the white matter at baseline (r = 0.515, B_10_ = 3.91), meaning that individuals with high plasma GDF15 measures across timepoints tended to exhibit high MitoD in the white matter of the brain.

**Figure 5.**
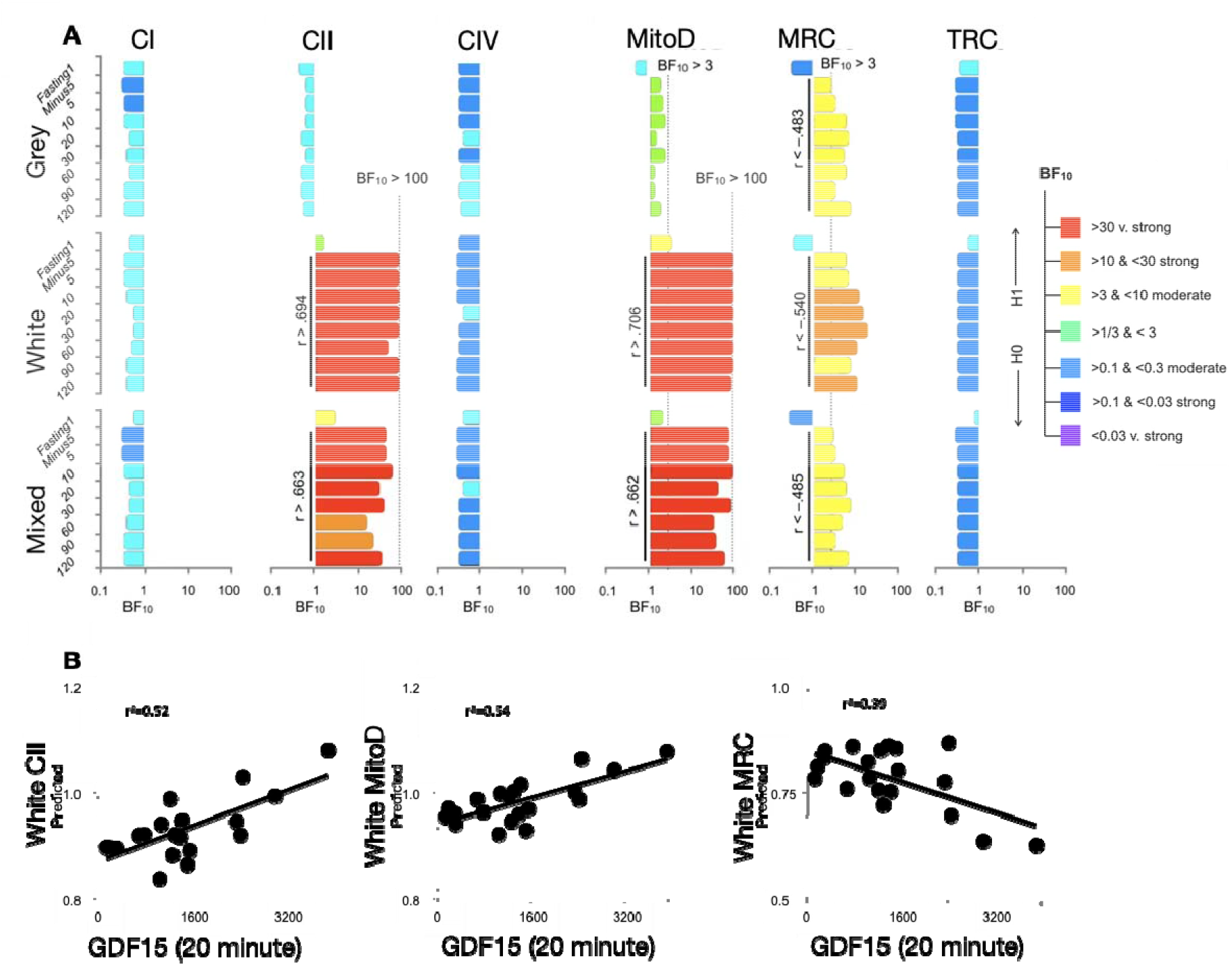
Associations between mitochondrial estimates and plasma GDF15. **(A**) Bayes factor (BF_10_) profiles showing the strength of association between plasma GDF15 and predicted mitochondrial features before and after a stress task. Strong to extreme evidence (BF_10_ > 30-100) was observed for positive relationships between GDF15 and complex II (CII) and mitochondrial density (MitoD), particularly in white and mixed tissue. In contrast, moderate evidence supported negative associations with mitochondrial respiratory capacity (MRC). (**B**) Scatterplots of the relationship between GDF15 20 minutes after a stress task and CII, mitochondrial density and MRC in the white matter.

More robust relationships existed around the stress manipulation, with strong to extreme evidence that elevated GDF15 levels from 5 minutes before and up to 2 hours after the stress tasks were related to higher mitochondrial density (white matter: r > 0.71, B_10_ > 141.77; mixed tissue: r > 0.66, B_10_ > 39.57) and increased Complex II expression (white matter: r > 0.69, B_10_ > 58.50; mixed tissues: r > 0.63, B_10_ > 18.21), in line again with a compensatory upregulation of mitochondrial content (both MitoD and CII are upregulated in OxPhos defects).

Additionally, post-stress GDF15 levels were moderately associated with reduced MRC across tissue types (grey matter: r < −0.48, B_10_ > 3.02; white matter: r < −0.54, B_10_ > 6.22; mixed tissues: r < −0.48, B_10_ > 3.1), consistent with the pathophysiology of mtDNA defects expected to specifically decrease MRC (the quality of each mitochondrion).

**Supplementary Table 3.**
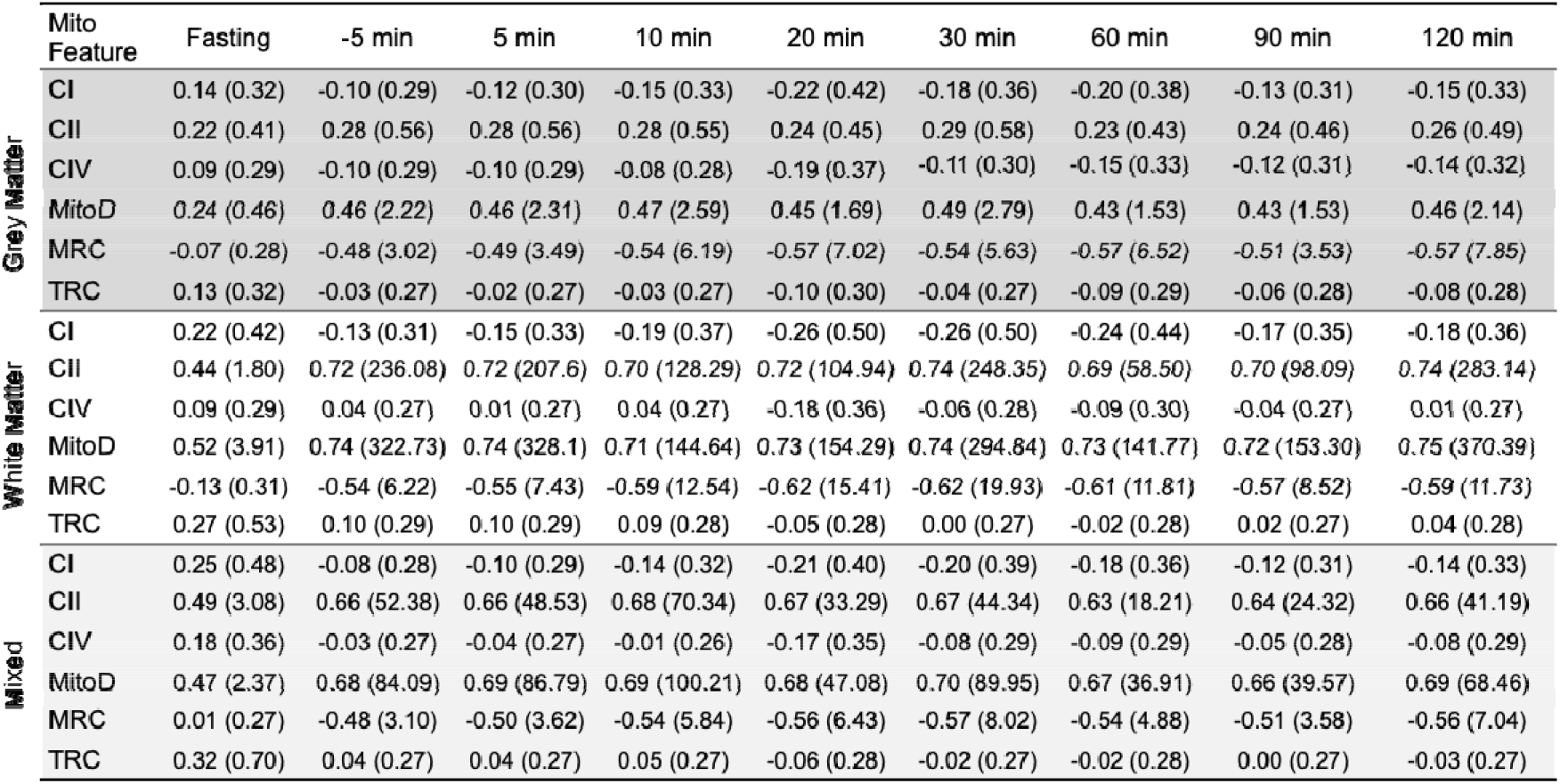
Predicted mitochondrial features and their relationship with plasma GDF15 at multiple time points surrounding the stress task. Pearson r values are shown in each cell, with corresponding Bayes factor (BF_10_) in brackets.

### Brain-behavior associations

Because mitochondrial efficiency is expected to support brain function, particularly neuronal function, we explored the relationship between predicted mitochondrial features in patients with mitochondrial genetic defects and a battery of 32 cognitive measures. An initial analysis identified several associations between neuropsychological scores and mitochondrial values (**Figure 6A**), although did not survive correction for multiple comparisons. Given the substantial intercorrelation among cognitive tests^38, 39^, we adopted a dimensionality-reduction approach to capture shared variance across measures while limiting the multiple-testing burden. Missing values (2.6%) were linearly interpolated^40^, and we used a varimax-rotated principal component analysis (PCA) on the full cognitive battery, resulting in 10 principal components that, together, explained ~80% of the variance in patients with mitochondrial diseases (see **Supplementary Table 4** for standardized component loadings).

**Figure 6.**
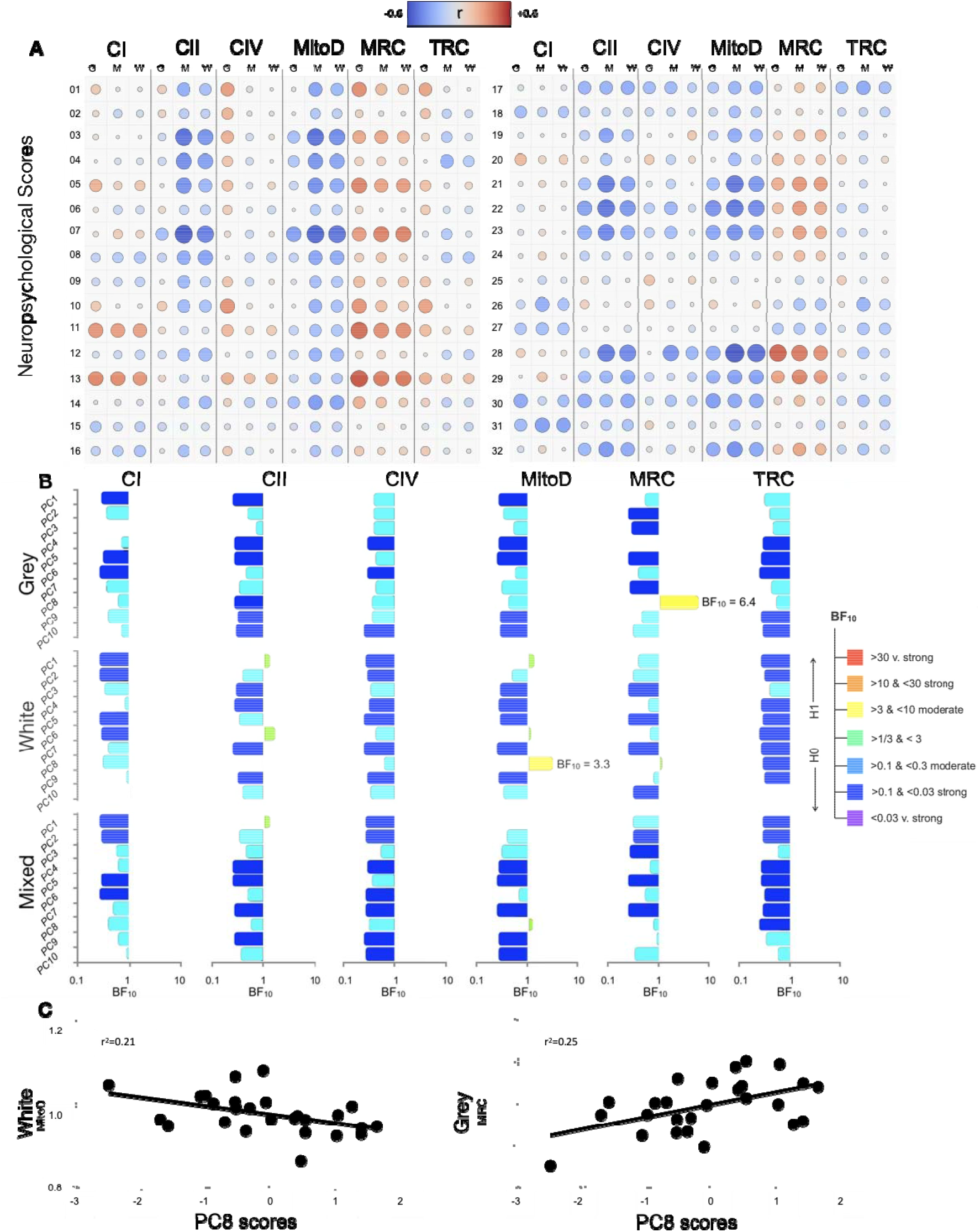
| Associations between mitochondrial estimates and cognitive performance. (**A**) Simple bivariate Pearson correlation between mitochondrial features and neuropsychological scores. (1) NAB Screening Numbers & Letters Part A Speed, (2) Color-Word Interference Condition 4: Inhibition/Switching, (3) Trail Making Condition 3: Letter Sequencing, (4) Trail Making Condition 2: Number Sequencing, (5) RBANS Total Score, (6) Color-Word Interference Condition 3: Inhibition, (7) Trail Making Condition 4: Number-Letter Switching, (8) Color-Word Interference Condition 2: Word Reading, (9) NAB Screening Numbers & Letters Part B Speed, (10) NAB Screening Numbers & Letters Part A Efficiency, (11) NAB Screening Digit Forward Longest Span, (12) Color-Word Interference Condition 1: Color Naming, (13) NAB Screening Digit Forward, (14) NAB Screening Shape Learning Delayed Recognition, (15) Verbal Fluency Condition 1: Category Switching Total Correct, (16) NAB Screening Numbers & Letters Part B Efficiency, (17) RBANS Picture Naming Total Score, (18) Verbal Fluency Condition 1: Category Switching Total Accuracy, (19) Verbal Fluency Condition 1: Category Fluency Total Correct, (20) RBANS List Learning Total Score, (21) Verbal Fluency Condition 1: Letter Fluency Total Correct, (22) NAB Screening Digit Backward, (23) NAB Screening Digit Backward Longest Span, (24) WASI Vocabulary Raw Score, (25) NAB Screening Numbers & Letters Part A Errors, (26) NAB Screening Numbers & Letters Part B Errors, (27) NAB Screening Shape Learning Percent Retention, (28) NAB Screening Shape Learning Immediate Recognition, (29) TOPF Total Score, (30) WASI Matrix Reasoning Raw Score, (31) RBANS Line Orientation Total Score, (32) Confirmed Correct Sorts. (**B**) Bayesian correlations between mitochondrial features and cognitive principal components (PC1-PC10, Supplementary Table 4). Moderate evidence of a relationship was found between PC8, reflecting accuracy and error monitoring, and mitochondrial density (white matter) and respiratory capacity (grey matter). (**C**) Scatterplots of the relationship between PC8 scores and mitochondrial density in the white matter and MRC in the grey matter.

These components appeared interpretable with PC1 corresponding to processing speed and executive throughput, PC2 to cognitive flexibility, PC3 to visuospatial reasoning, PC4 to working memory capacity, PC5 to lexical semantic retrieval, PC6 to learning encoding efficiency, PC7 to attention and inhibition, PC8 to accuracy and error monitoring, and PC9 to visual memory. PC10 explained minimal variance related to a single neuropsychological score (i.e., NAB screening numbers and letters part B errors). **Figure 6** shows that a moderate relationship exists between lowerPC8 scores (worse performance) and increased mitochondrial density (r = −0.456, B_10_ = 3.26) and decreased mitochondrial respiratory capacity (r = 0.504, B_10_ = 6.41). These data buttress the conclusion that cognitive function is more severely affected in patients with lower MRC (due to individual mitochondria inefficiency) and higher MitoD (due to compensatory upregulation of mitogenesis).

**Supplementary Table 4.** List of neuropsychological scores and their squared standardized principal component analysis loadings. Rows are colored/bold according to their dominant principal component.

| Neuropsychological Scores/PCs | PC1 | PC2 | PC3 | PC4 | PC5 | PC6 | PC7 | PC8 | PC9 | PC10 |
| --- | --- | --- | --- | --- | --- | --- | --- | --- | --- | --- |
| Screening Numbers & Letters Part A Efficiency | **0.64** | 0.02 | 0.01 | 0.01 | 0.00 | 0.00 | 0.04 | 0.07 | 0.02 | 0.01 |
| Screening Numbers & Letters Part A Speed | **0.63** | 0.00 | 0.00 | 0.01 | 0.00 | 0.00 | 0.09 | 0.11 | 0.02 | 0.02 |
| Condition1: Color Naming | **0.57** | 0.08 | 0.01 | 0.04 | 0.05 | 0.01 | 0.02 | 0.01 | 0.00 | 0.00 |
| Condition2: Word Reading | **0.56** | 0.10 | 0.00 | 0.07 | 0.05 | 0.00 | 0.01 | 0.00 | 0.01 | 0.00 |
| RBANS Total Score | **0.49** | 0.07 | 0.00 | 0.01 | 0.04 | 0.04 | 0.00 | 0.05 | 0.00 | 0.00 |
| Condition3: Inhibition | **0.45** | 0.02 | 0.12 | 0.04 | 0.08 | 0.01 | 0.00 | 0.02 | 0.00 | 0.00 |
| Condition3: Letter Sequencing | **0.40** | 0.08 | 0.01 | 0.01 | 0.20 | 0.01 | 0.01 | 0.00 | 0.04 | 0.11 |
| Condition4: Number-Letter Switching | **0.29** | 0.08 | 0.19 | 0.03 | 0.16 | 0.02 | 0.02 | 0.00 | 0.01 | 0.02 |
| Condition 1: Category Switching Total Switching Accuracy | 0.00 | **0.73** | 0.04 | 0.00 | 0.00 | 0.00 | 0.03 | 0.01 | 0.02 | 0.00 |
| Condition 1: Category Switching Total Correct | 0.01 | **0.69** | 0.07 | 0.00 | 0.00 | 0.00 | 0.04 | 0.01 | 0.04 | 0.00 |
| Condition 1: Category Fluency Total Correct | 0.08 | **0.62** | 0.00 | 0.02 | 0.04 | 0.01 | 0.01 | 0.00 | 0.01 | 0.00 |
| Condition 1: Letter Fluency Total Correct | 0.11 | **0.39** | 0.01 | 0.03 | 0.03 | 0.08 | 0.02 | 0.00 | 0.00 | 0.01 |
| RBANS List Learning Total Score | 0.02 | 0.15 | **0.02** | 0.03 | 0.14 | 0.10 | 0.01 | 0.08 | 0.00 | 0.08 |
| Matrix Reasoning Raw Score | 0.01 | 0.00 | **0.71** | 0.00 | 0.00 | 0.06 | 0.02 | 0.00 | 0.01 | 0.00 |
| Confirmed Correct Sorts | 0.02 | 0.06 | **0.56** | 0.00 | 0.06 | 0.02 | 0.02 | 0.02 | 0.01 | 0.00 |
| RBANS Line Orientation Total Score | 0.00 | 0.00 | **0.47** | 0.00 | 0.01 | 0.04 | 0.00 | 0.03 | 0.02 | 0.04 |
| Vocabulary raw score | 0.01 | 0.14 | **0.34** | 0.18 | 0.03 | 0.00 | 0.00 | 0.03 | 0.02 | 0.05 |
| Screening Digital Forward Longest Span | 0.02 | 0.00 | 0.00 | **0.84** | 0.01 | 0.03 | 0.00 | 0.00 | 0.01 | 0.00 |
| Screening Digital Forward | 0.02 | 0.00 | 0.00 | **0.82** | 0.01 | 0.06 | 0.00 | 0.00 | 0.00 | 0.00 |
| TOPF Total | 0.07 | 0.17 | 0.21 | **0.24** | 0.00 | 0.01 | 0.01 | 0.00 | 0.06 | 0.00 |
| RBANS Picture Naming Total Score | 0.07 | 0.06 | 0.10 | **0.23** | 0.01 | 0.01 | 0.02 | 0.01 | 0.02 | 0.00 |
| Screening Numbers & Letters Part A Error | 0.01 | 0.00 | 0.00 | 0.00 | **0.70** | 0.00 | 0.02 | 0.00 | 0.02 | 0.00 |
| Condition2: Number Sequencing | 0.14 | 0.07 | 0.01 | 0.03 | **0.32** | 0.00 | 0.02 | 0.05 | 0.05 | 0.02 |
| Screening Digital Backward Longest Span | 0.02 | 0.01 | 0.06 | 0.03 | 0.01 | **0.81** | 0.00 | 0.01 | 0.00 | 0.00 |
| Screening Digital Backward | 0.02 | 0.00 | 0.07 | 0.04 | 0.00 | **0.80** | 0.00 | 0.00 | 0.00 | 0.00 |
| Screening Numbers & Letters Part B Efficiency | 0.04 | 0.02 | 0.02 | 0.00 | 0.01 | 0.00 | **0.81** | 0.00 | 0.00 | 0.01 |
| NAB screening numbers and letters part b speed | 0.10 | 0.04 | 0.01 | 0.00 | 0.05 | 0.00 | **0.68** | 0.01 | 0.01 | 0.00 |
| Screening Shape Learning Immediate Recognition | 0.04 | 0.01 | 0.03 | 0.00 | 0.00 | 0.00 | 0.01 | **0.72** | 0.10 | 0.00 |
| Screening Shape Learning Delayed Recognition | 0.04 | 0.01 | 0.04 | 0.00 | 0.00 | 0.01 | 0.00 | **0.51** | 0.15 | 0.03 |
| Screening Shape Learning Percent Retention | 0.00 | 0.01 | 0.01 | 0.00 | 0.02 | 0.01 | 0.02 | 0.01 | **0.77** | 0.02 |
| Condition4: Inhibition/Switching | 0.07 | 0.02 | 0.03 | 0.09 | 0.11 | 0.01 | 0.02 | 0.05 | **0.22** | 0.11 |
| NAB screening numbers and letters part b errors | 0.00 | 0.00 | 0.00 | 0.01 | 0.01 | 0.00 | 0.01 | 0.02 | 0.01 | **0.67** |

## Discussion

Our findings show that mitochondrial phenotypes reconstructed from multimodal neuroimaging capture biologically meaningful variation across age, genetic mitochondrial defects, systemic energetic stress and cognitive functions.

Perhaps the most intriguing finding from this work is that component-specific mitochondrial biochemical signatures are detectable by MRI. The agreement between the expected hallmark signature (elevated complex II and mitoD as compensation for impaired complexes I and IV) of mitochondrial diseases and our observations implies that the MR signal modalities contain information not only about mitochondrial abundance and health in general, but about specific mitochondrial components (i.e., specific enzymes). One possibility for this observation is the varying amount and quality of paramagnetic iron in each of the respiratory chain enzymes (8 iron-sulfur clusters in complex I, 3 iron-sulfur clusters in complex II, 2 iron-copper containing heme molecules in complex IV)^41, 42^. The stochiometry of iron and sulfur atoms also varies among clusters in different complexes. Another possible explanation for our results is that the strong transmembrane proton gradient across the inner mitochondrial membrane^43-45^ contributes relevant information detected by general proton-based MR modality. Complexes I and IV contribute to the large proton concentration (localized low pH) within well-functioning mitochondria. This could, in theory, account for why T1- and T2-weighted brain imaging, which are sensitive to proton relaxation properties,) would include mitochondria-relevant data. Further work will be necessary to rigorously probe the physical and biochemical basis of these findings.

Biologically, one of our findings most consistent with extensive cellular^46^ and animal^47, 48^ literature is the robust age-related decline in mitochondrial density and tissue respiratory capacity. In comparison to the change in mitochondrial content, intrinsic mitochondrial respiratory capacity remained relatively preserved through late midlife (i.e., 60 years old), as demonstrated for the relative preservation of the respiratory chain with age in animal models^49^. This pattern confirms that ageing is primarily associated with a reduction in mitochondrial abundance and turnover, rather than a loss of intrinsic respiratory efficiency of the remaining organelles–a notion recently confirmed in aging human skeletal muscle across the adult lifespan^22, 50^.

Mitochondrial genetic defects are expected to undermine the ability of individual mitochondria to generate ATP, which directly elicit compensatory mechanisms. These include increased mitochondrial content and preferential reliance on nuclear-encoded respiratory chain components. Specifically, complex II (made of 4 subunits) is entirely nuclear-encoded^51^, and is therefore not biochemically affected by mitochondrial DNA mutations. As a result, while respiratory chain complexes I and IV are defective in mtDNA diseases, complex II often exhibits elevated activity, due to compensatory upregulation driven by mitochondrial biogenesis^30^. In line with this framework, our group-level analyses revealed increased complex II activity and mitochondrial density in patients with mitochondrial diseases. Together, these changes contributed to a preserved or an apparent increase in tissue respiratory capacity (TRC) particularly affecting the striatal, fronto-parietal, orbitofrontal, and occipital regions.

Beyond these quantitative changes, our correlation and Bayesian analyses revealed marked alterations in the coupling among mitochondrial features in patients. This reflects the expected disruption of mitochondrial homeostasis in people with mitochondrial disease. Whereas mitochondrial features were tightly coordinated in healthy individuals, patients exhibited a reconfiguration of mitochondrial features’ relationships, consistent with the biochemical effects of mtDNA gene defects. Such reconfiguration aligns with models of compensatory mitochondrial proliferation and oxidative phosphorylation rebalancing aimed at sustaining energetic output in spite of impaired mitochondrial energy transformation capacity^52^. In contrast, in our very small sample of MELAS patients with sufficient neuroimaging coverage (n=2), compensatory mechanisms appear to fail, as previously described^30^, leading to reduced mitochondrial content and no increase in complex II. Overall, these results suggest that our prediction model can capture compensatory mitochondrial proliferation and homeostatic strategies to sustain oxidative phosphorylation despite impaired mitochondrial transcription.

As a final piece of evidence that our MR-based predicted mitochondrial features indeed tap into mitochondrial biology, we observed an association between predicted measures of mitochondrial features and symptoms consistent with that classically reported in patients with mitochondrial diseases. In particular, the compensatory complex II activity and increased mitochondrial density were strongly associated with the biomarker of energetic stress GDF15. Moreover, the magnitude of this association was stronger when blood GDF15 was measured during psychological stress compared to when GDF15 was measured the same day, but under resting conditions. This pattern supports the existence of a bioenergetic-stress coupling mechanism whereby mitochondrial adaptations to energetic demand are translated into a metabolic signal ^53^, here GDF15. In parallel, increased mitochondrial density in the white matter and reduced mitochondrial respiratory capacity in the grey matter were associated with poorer performance in accuracy in error monitoring, consistent with the neuropsychological profile of mitochondrial diseases^54^. These brain mitochondria-human behavior findings suggest that mitochondrial disease lies along a continuum in which compensatory bioenergetic response, while supporting baseline function, tracks with physiological stress signaling and cognitive costs.

Mitochondrial neuroimaging is relatively new and, accordingly, not exempt from limitations that warrant acknowledgement and further investigation. For instance, mitochondrial mapping is anchored to the biochemical measurements from a single post-mortem coronal section of one brain as no individual-level mitochondrial ground truth is possible for any living participants. In spite of this limitation, every external criterion we examined (age, genotype, GDF15 or cognition) was concordant with the literature. Additionally, the disease group is rare and, accordingly, small and limited in power. This precludes any formal inference for the MELAS group but significant results underline the sensitivity of some of our mitochondrial measures to disease. Finally, these findings are derived from a single cohort and should be replicated, ideally against orthogonal in vivo measures of brain energetics^55-57^.

Overall, we provide initial evidence that neuroimaging-derived mitochondrial phenotypes capture meaningful biological variation across age, mitochondrial homeostasis, genetics, stress physiology, and cognition, supporting their future use as a non-invasive window onto human brain bioenergetics. Importantly, these effects cannot be attributed to any single neuroimaging modality, as mitochondrial features showed stronger Bayesian evidence than their constituent MRI measures and exhibited biologically selective dissociations inconsistent with nonspecific MRI-driven effects. Together, these findings call for further validation work examining neuroimaging-based mitochondrial mapping as a non-invasive tool to probe mitochondrial biology and compensatory responses, disease burden, and potential treatment response in the living human brain.

## Online Methods

### Participants

Participants were drawn from the Mitochondrial Stress, Brain Imaging, and Epigenetics (MiSBIE) study, carried out at Columbia University Irving Medical Center under the oversight of the New York State Psychiatric Institute Institutional Review Board (protocol #7424) and registered on ClinicalTrials.gov (NCT04831424). Recruitment occurred between 2018 and 2023 and targeted English-speaking adults aged 18-60 years. Volunteers were enrolled via clinical and research pathways at Columbia University (including the Neuromuscular Clinic and RecruitMe), the Children’s Hospital of Philadelphia, and national/international mitochondrial disease organizations (e.g., NAMDC, UMDF). Individuals were not eligible if they had a recent acute illness, were pregnant, had an active cancer diagnosis, were using systemic steroids, were participating in another clinical trial, or showed evidence of cognitive impairment on the Telephone Interview for Cognitive Status (TICS-41 inferior or equal to 30).

The protocol consisted of two consecutive study days. Day 1 included biospecimen collection, clinical examination, standardized stress reactivity and recovery session, detailed physiological phenotyping, functional/standing capacity assessment, resting metabolic rate measurement, psychosocial questionnaires, and neuropsychological testing. Day 2 comprised MRI at the Zuckerman Institute, including high-resolution structural imaging (T1-and T2-weighted scans), field mapping, resting-state fMRI, diffusion-weighted imaging, and task-based fMRI.

Across the parent MiSBIE cohort, 110 participants were enrolled (40 with mitochondrial disease and 70 healthy controls). However, the present analyses were restricted to the subset with complete neuroimaging, as this was needed to estimate the mitochondrial measures required for this manuscript. This yielded a final sample of 85 participants. Demographics of the population studied in the present paper are reported in Table 1.

**Table 1.** Demographic and clinical characteristics of participants included in the study. Clinical severity was assessed using the Newcastle Mitochondrial Disease Adult Scale (NMDAS).

| Group | N | Female, n (%) | Age, Mean ± SD (years) | NMDAS |
| --- | --- | --- | --- | --- |
| Controls | 59 | 41 (69.5%) | 37.2 ± 10.7 | 2.23 ± 3.12 |
| Single Deletion | 14 | 11 (78.6%) | 41.4 ± 7.6 | 4.4 ± 3.9 |
| 3243 A>G | 10 | 6 (60%) | 32.1 ± 8.6 | 14 ± 14.1 |
| MELAS | 2 | 1 (50%) | 43.1 ± 10.8 | 20.6 ± 11.21 |
| Mito Disease | 26 | 18 (69.2%) | 37.1 ± 9.8 | 11.6 ± 11.1 |
| Total | 85 | 59 (69.4%) | 37.1 ± 10.4 | 4.9 ± 7.7 |

### Neuroimaging Acquisitions

Magnetic resonance imaging data were acquired on a 3T Magnetom Aera with a Head/Neck 64-channel coil.

T1-weighted images were obtained using a three-dimensional sequence with the following parameters: repetition time (TR) = 2.3 s, echo time (TE) = 3.49 ms, flip angle = 8°, voxel size = 1x1x1mm with slice thickness of 1 mm. Parallel imaging was applied with an acceleration factor of 2.

T2-weighted images were acquired with a TR = 3.2s, TE = 562 ms, flip angle = 120°, a voxel size of 0.7 x 0.7 x 0.7 mm, and parallel imaging with an acceleration factor of 2.

Diffusion-weighted imaging (DWI) was acquired using a two-dimensional axial EPI sequence, producing a 120 x 120 x 80 matrix with voxel dimensions of 1.8 x 1.8 x 1.8 mm. Acquisitions were performed using a TR of 3540ms and a TE of 75 ms, a multiband acceleration factor of 3, no in-plane parallel acceleration (R = 1), partial Fourier factor of 6/8, and fat saturation. Diffusion was measured along the anterior-posterior phase of acquisition and multiple gradients using four different B-values (8 repetitions x B = 0 s.mm^−2^; 8 directions x B = 300 s.mm^−2^; 30 directions x B = 1000 s.mm^−2^; 60 directions x B = 2000 s.mm^−2^). In order to calculate the correction for susceptibility artefact, an additional 8 repetitions x B = 0 s.mm^−2^ was acquired along the posterior-anterior phase of acquisition.

Functional data were collected on the same scanner to estimate blood-oxygen-level-dependent (BOLD) signal fluctuations during rest. Whole-brain echo-planar imaging (EPI) was performed in the axial plane using multiband acceleration (=8) with a TR of 460 ms and a TE of 29 ms, a flip angle = 44°, and fat suppression with a resolution of 3.02 x 3.02 x 3 mm.

### Neuroimaging Processing

Structural MRI data were processed to derive complementary measures of cortical morphology and tissue composition. All these steps were performed in the native space. Cortical thickness maps were derived from T1-weighted images using the cortical thickness pipeline (Diffeomorphic Registration based Cortical Thickness, DiReCT^58^) implemented in Advanced Normalisation Tool (ANTs^59^) and made available via BCBtoolkit (www.bcblab.com^60^).

Voxelwise probability maps of grey matter and white matter were generated from T1-weighted images using the unified segmentation framework^61^ implemented in Statistical Parametric Mapping (SPM).

To estimate T1w/T2w^62, 63^, T1-weighted and T2-weighted images were first coregistered using ANTs, and a voxelwise T1w/T2w ratio map was computed. This ratio has been argued to enhance contrast related to microstructure.

Diffusion-weighted imaging was preprocessed using the Human Connectome Project (HCP) diffusion pipeline^64^, following established procedures for correction of susceptibility and eddy-current artefacts, as well as subject motion. The susceptibility-induced artefacts were estimated by exploiting the pairs of B0 images acquired with opposite phase-encoding directions^65^. This estimation was then used to correct geometric distortion across the full dataset using TOPUP in the FMRIB Software Library (FSL^66^). Next, residual head motion and eddy current distortions were corrected using EDDY as part of FSL. Corrected data were subsequently entered into StarTrack (https://www.mr-startrack.com/) for diffusion tensor imaging map estimation (fractional anisotropy, FA; radial diffusivity, RD; axial diffusivity, AD; mean diffusivity, MD) and tractography. For the tractography, fibre orientation distributions (FOD) were estimated using damped Richardson-Lucy spherical deconvolution algorithm^67, 68^. A fixed single fiber response function was assumed, corresponding to a shape factor α = 1.5 x 10^−3^mm^2^.s^−1^, together with a geometric damping parameter set to 8. Spherical deconvolutions were iterated 200 times to ensure stable convergence of the FOD estimates. To suppress spurious orientations, an absolute amplitude threshold was set to 3 times the FOD amplitude measured in a grey voxel. A relative threshold of 8% of the maximum FOD peak was also applied. Whole-brain streamline tractography was then performed using a modified Euler algorithm. Streamlines were propagated with a step size of 0.5mm, a maximum turning angle of 35°, and a minimum streamline length of 15mm. Streamline density maps were derived from whole-brain tractography using trackvis (https://trackvis.org). Additional diffusion metrics such as intra-cellular volume fraction (ICVF) extra-cellular volume fraction (ISOCF), and orientation dispersion index (i.e. neurite complexity, OD) were estimated using the Accelerated Microstructure Imaging Via Convex Optimization (AMICO) framework^69^.

Finally, resting-state functional MRI data were processed using fMRIprep^70^. Following preprocessing, the resting-state data underwent additional denoising steps, including nuisance regressors (24 head-motion parameters) and the cerebrospinal fluid signal and its derivatives. Spike regressors derived from Mahalanobis distance ^71^ were employed to mitigate the transient artefact. Specifically, volumes whose distance exceeded 95% confidence intervals of the multivariate distribution were discarded. Each identified outlier volume was modeled with a binary regressor. The denoised time series were subsequently detrended and spatially smoothed using a Gaussian kernel with full half-width maximum = 4mm. Regional homogeneity (ReHo) and amplitude of low-frequency fluctuations (ALFF) as well as the ratio between low and high frequency fluctuations (fALFF) were calculated using the Data Processing & Analysis Imaging (DPABI) toolbox^72^. For ALFF, a temporal band pass filter was applied in the range of 0.01-0.10 hertz, while for ReHo, we used a 0.01-0.08 hertz filter. Additionally, the maximum level of activity was extracted for each voxel using the fslmaths function in FSL. Finally, Shannon entropy maps were extracted from the previously preprocessed rs-fMRI using the following formula: − sum (p*log(p)), where p indicates the probability of the intensity in the voxel^73^.

### Templates

10 healthy MiSBIE participants who most closely matched the MitoBrainMap post-mortem brain (54-year-old neurotypical male) in sex and age (males, 36-54 years) were used to generate diffusion, structural, and fMRI templates using the ANTs multitemplate construction framework. Images derived from the same source modality were grouped prior to template generation (i.e., structural, diffusion and functional). This modality-specific approach avoids cross-modality bias during template estimation and ensures optimal anatomical correspondence within each image class. Template construction was performed using the ANTs multitemplate iterative procedure, which alternates iteratively between averaging and diffeomorphic registration to estimate a representative population template for each modality (see **supplementary figure 1C**). For simplicity, the resulting modality-specific templates were linearly registered to the MNI space.

### Neuropsychological Assessments

Neuropsychological tests were administered by the study coordinator, who was trained and tested by a neuropsychologist, in accordance with test-specific requirements and specifications. The test battery was delivered virtually from 2021 onwards due to the COVID-19 pandemic^17^.

Briefly, the Test of Premorbid Functioning (TOPF, Pearson Assessments, 015800972X, Coushatta,LA) was used to assess individuals’ cognitive and memory function. The Wechsler Abbreviated Scale of Intelligence Second Edition (WASI-II, Pearson Assessments, 0158981561, Coushatta, LA) was used to assess general intellectual ability. The Repeatable Battery for the Assessment of Neuropsychological Status (RBANS, PearsonAssessments, 0158006976, Coushatta, LA) was used to assess visuospatial judgement, confrontation naming, verbal learning and memory, and processing speed. The Neuropsychological Assessment Battery (NAB, Psychological Assessment Resources,11918-KD, Lutz, FL) was used to assess visual learning and memory, attention and inhibition, and attention and working memory. The Delis-Kaplan Executive Function System test (DKEFS, Pearson Assessments,0158091108, Coushatta, LA) was used to assess letter and category fluency, verbal set-shifting, working memory, cognitive flexibility and speed, response inhibition and mental flexibility and conceptualization.

### Blood Collection and GDF15 Measurements

GDF15 was measured in plasma via a central venous catheter at 9 time points^17^. Briefly, one sample was collected in the morning in a fasted state, while eight samples were collected in the afternoon across the stress reactivity protocol involving a socio-evaluative speech task where participants were asked to defend themselves against an alleged transgression (shoplifting). 16-20 mL of whole blood was collected at the morning fasting timepoint in two 10 mL K_2_EDTA blood collection tubes (BD #366643). 5 mL of whole blood was collected at each afternoon time point in 6 mL K_2_EDTA blood collection tubes (BD #367899).

Immediately after collection, whole blood was inverted 10–12 times and centrifuged at 2000g for 3.5 min at room temperature. Samples were then placed on ice and transported to the laboratory, where samples were immediately centrifuged at 2000g for 10min at 4 °C. Approximately 80% of the plasma was collected from the upper portion of each tube and transferred into new 15 mL conical tubes. Tubes were then centrifuged at 2000g for another 10min at 4 °C. Approximately 90% of the plasma supernatant was then transferred to new 15 mL conical tubes and inverted to mix. 0.5– 1.5 mL aliquots per timepoint were made with the resulting cell-free plasma, and stored at −80 °C before GDF15 measurement.

Plasma GDF15 levels were quantified using a high-sensitivity ELISA kit (R&D Systems, DGD150, SGD150) following the manufacturer’s instructions. The detailed measurement protocol can be found at^16^. Briefly, plasma was mixed with assay diluent at a 1:4 ratio. Plates were read at an absorbance nm at 450 nm on a Spectramax M2 (Spectramax Pro 6, Molecular Devices), and concentrations were interpolated using a four-parameter logistic (4PL) model in GraphPad Prism (version 10.3.1) using the standard curve (five concentration points) run on each plate. Two or three reference samples were run per plate and their inter-assay coefficients of variation (CV) were monitored. All samples for a particular individual were kept on the same plate where possible. Each unique participant x timepoint sample was run in duplicate on separate plates where possible, and final concentrations were calculated using an average of the duplicates. If duplicates exhibited a C.V. larger than 15%, they were rerun if sample volume allowed. Values below the minimum detection range (2.0 pg/ml) were considered undetectable and excluded from analyses.

## Statistics

Hierarchical regression with backward elimination and principal component analyses were performed in SPSS (https://www.ibm.com/products/spss-statistics), Bayesian statistics in JASP (https://jasp-stats.org/)

## Data and Code Availability Statement

The MiSBIE study protocol is available and data can be requested at www.picardlab.com/MiSBIE.

## Author Contributions

M.T.d.S., E.V.M. and M.P. designed the study, interpreted the results and drafted the manuscript. M.T.d.S. developed the MitoBrain model. M.T.d.S., E.V.M., CCL & KB performed data analyses. CCL and CK collected data. M.T.d.S., E.V.M. and CCL prepared the figures. T.D.W. revised the manuscript. M.P. supervised the parent MiSBIE study and analyses. All authors reviewed the final version of the manuscript.

## REFERENCES

1. Sagan, L. On the origin of mitosing cells. Journal of Theoretical Biology 14, 225–IN226 (1967).

2. Monzel, A.S., Enríquez, J.A. & Picard, M. Multifaceted mitochondria: moving mitochondrial science beyond function and dysfunction. Nat Metab 5, 546–562 (2023).

3. Lane, N. & Martin, W. The energetics of genome complexity. Nature 467, 929–934 (2010).

4. Picard, M. & Shirihai, O.S. Mitochondrial signal transduction. Cell Metab 34, 1620–1653 (2022).

5. Theriault, J.E., et al. It’s not the thought that counts: Allostasis at the core of brain function. Neuron 113, 4107–4133 (2025).

6. Kann, O. & Kovács, R. Mitochondria and neuronal activity. American Journal of Physiology-Cell Physiology 292, C641–C657 (2007).

7. Mattson, M.P., Gleichmann, M. & Cheng, A. Mitochondria in neuroplasticity and neurological disorders. Neuron 60, 748–766 (2008).

8. Iwata, R. & Vanderhaeghen, P. Regulatory roles of mitochondria and metabolism in neurogenesis. Current Opinion in Neurobiology 69, 231–240 (2021).

9. Hollis, F., et al. Mitochondrial function in the brain links anxiety with social subordination. Proceedings of the National Academy of Sciences 112, 15486–15491 (2015).

10. Rosenberg, A.M., et al. Brain mitochondrial diversity and network organization predict anxiety-like behavior in male mice. Nature Communications 14, 4726 (2023).

11. Kaufmann, P., et al. Natural history of MELAS associated with mitochondrial DNA m.3243A>G genotype. Neurology 77, 1965–1971 (2011).

12. Shatalina, E., et al. Mitochondria and Cognition: An [18F]BCPP-EF Positron Emission Tomography Study of Mitochondrial Complex I Levels and Brain Activation During Task Switching. Biological Psychiatry: Cognitive Neuroscience and Neuroimaging 10, 823–832 (2025).

13. Mosharov, E.V., et al. A human brain map of mitochondrial respiratory capacity and diversity. Nature (2025).

14. Gorman, G.S., et al. Mitochondrial diseases. Nature Reviews Disease Primers 2, 16080 (2016).

15. Lin, Y., et al. Accuracy of FGF-21 and GDF-15 for the diagnosis of mitochondrial disorders: A meta-analysis. Ann Clin Transl Neurol 7, 1204–1213 (2020).

16. Huang, Q., et al. The mitochondrial disease biomarker GDF15 is dynamic, quantifiable in saliva, and correlates with disease severity. Molecular Genetics and Metabolism 145, 109179 (2025).

17. Kelly, C., et al. A platform to map the mind-mitochondria connection and the hallmarks of psychobiology: the MiSBIE study. Trends Endocrinol Metab 35, 884–901 (2024).

18. Tsuchida, A., et al. The MRi-Share database: brain imaging in a cross-sectional cohort of 1870 university students. Brain Struct Funct 226, 2057–2085 (2021).

19. Torii, K., Sugiyama, S., Takagi, K., Satake, T. & Ozawa, T. Age-related Decrease in Respiratory Muscle Mitochondrial Function in Rats. American Journal of Respiratory Cell and Molecular Biology 6, 88–92 (1992).

20. Bratic, A. & Larsson, N.G. The role of mitochondria in aging. J Clin Invest 123, 951–957 (2013).

21. Omidsalar, A.A., et al. Mitochondrial DNA Variation in the Aging Human Cerebral Cortex and Cerebellum. Aging Cell 25, e70340 (2026).

22. Cefis, M., et al. Impact of physical activity on physical function, mitochondrial energetics, ROS production, and Ca(2+) handling across the adult lifespan in men. Cell Rep Med 6, 101968 (2025).

23. Keysers, C., Gazzola, V. & Wagenmakers, E.-J. Using Bayes factor hypothesis testing in neuroscience to establish evidence of absence. Nature Neuroscience 23, 788–799 (2020).

24. Howell, D.C. Statistical Methods for Psychology (Wadsworth Cengage Learning, Belmont, CA, 2013).

25. Vincent, A.E. & Picard, M. Multilevel heterogeneity of mitochondrial respiratory chain deficiency. J Pathol 246, 261–265 (2018).

26. Trifunov, S., et al. Clonal expansion of mtDNA deletions: different disease models assessed by digital droplet PCR in single muscle cells. Sci Rep 8, 11682 (2018).

27. Sitarz, K.S., et al. MFN2 mutations cause compensatory mitochondrial DNA proliferation. Brain 135, e219, 211-213; author reply e220, 211-213 (2012).

28. Sercel, A.J., et al. Hypermetabolism and energetic constraints in mitochondrial disorders. Nat Metab 6, 192–195 (2024).

29. Vincent, A.E., et al. Subcellular origin of mitochondrial DNA deletions in human skeletal muscle. Ann Neurol 84, 289–301 (2018).

30. Giordano, C., et al. Efficient mitochondrial biogenesis drives incomplete penetrance in Leber’s hereditary optic neuropathy. Brain 137, 335–353 (2014).

31. Letts, J.A. & Sazanov, L.A. Clarifying the supercomplex: the higher-order organization of the mitochondrial electron transport chain. Nat Struct Mol Biol 24, 800–808 (2017).

32. Trumpff, C., et al. Mitochondrial respiratory chain protein co-regulation in the human brain. Heliyon 8, e09353 (2022).

33. DiMauro, S. & Schon Eric, A. Mitochondrial Respiratory-Chain Diseases. New England Journal of Medicine 348, 2656–2668.

34. Sciacco, M., Bonilla, E., Schon, E.A., DiMauro, S. & Moraes, C.T. Distribution of wild-type and common deletion forms of mtDNA in normal and respiration-deficient muscle fibers from patients with mitochondrial myopathy. Hum Mol Genet 3, 13–19 (1994).

35. Yarkoni, T., Poldrack, R.A., Nichols, T.E., Van Essen, D.C. & Wager, T.D. Large-scale automated synthesis of human functional neuroimaging data. Nat Methods 8, 665–670 (2011).

36. Gorman, G.S., et al. Perceived fatigue is highly prevalent and debilitating in patients with mitochondrial disease. Neuromuscul Disord 25, 563–566 (2015).

37. Kelly, C., et al. Perceived association of mood and symptom severity in adults with mitochondrial diseases. Mitochondrion 84, 102033 (2025).

38. Cona, G.F., et al. Archetypes in human behavior and their brain correlates: An evolutionary trade-off approach. bioRxiv (2018).

39. Smith, S.M., et al. A positive-negative mode of population covariation links brain connectivity, demographics and behavior. Nature Neuroscience 18, 1565–1567 (2015).

40. Little, R.J.A. & Rubin, D.B. Statistical Analysis with Missing Data (John Wiley & Sons, Inc., Hoboken, NJ, 2002).

41. Read, A.D., Bentley, R.E., Archer, S.L. & Dunham-Snary, K.J. Mitochondrial iron-sulfur clusters: Structure, function, and an emerging role in vascular biology. Redox Biol 47, 102164 (2021).

42. Babbitt, S.E., Sutherland, M.C., San Francisco, B., Mendez, D.L. & Kranz, R.G. Mitochondrial cytochrome c biogenesis: no longer an enigma. Trends Biochem Sci 40, 446–455 (2015).

43. Lane, N. Why Are Cells Powered by Proton Gradients? Nature Education 3, 18 (2010).

44. Santo-Domingo, J. & Demaurex, N. Perspectives on: SGP symposium on mitochondrial physiology and medicine: the renaissance of mitochondrial pH. J Gen Physiol 139, 415–423 (2012).

45. Mitchell, P. & Moyle, J. Estimation of Membrane Potential and pH Difference across the Cristae Membrane of Rat Liver Mitochondria. European Journal of Biochemistry 7, 471–484 (1969).

46. Tauchi, H. & Sato, T. Age changes in size and number of mitochondria of human hepatic cells. J Gerontol 23, 454–461 (1968).

47. Herbener, G.H. A morphometric study of age-dependent changes in mitochondrial population of mouse liver and heart. J Gerontol 31, 8–12 (1976).

48. Stocco, D.M. & Hutson, J.C. Quantitation of mitochondrial DNA and protein in the liver of Fischer 344 rats during aging. J Gerontol 33, 802–809 (1978).

49. Navarro, A. & Boveris, A. The mitochondrial energy transduction system and the aging process. Am J Physiol Cell Physiol 292, C670–686 (2007).

50. Short, K.R., et al. Decline in skeletal muscle mitochondrial function with aging in humans. Proc Natl Acad Sci U S A 102, 5618–5623 (2005).

51. Leonard, J.V. & Schapira, A.H.V. Mitochondrial respiratory chain disorders I: mitochondrial DNA defects. The Lancet 355, 299–304 (2000).

52. Wallace, D.C. Mitochondrial Diseases in Man and Mouse. Science 283, 1482–1488 (1999).

53. Picard, M. & McEwen, B.S. Psychological Stress and Mitochondria: A Systematic Review. Biopsychosocial Science and Medicine 80 (2018).

54. Finsterer, J. Cognitive dysfunction in mitochondrial disorders. Acta Neurologica Scandinavica 126, 1–11 (2012).

55. Li, Y., et al. Multiplexed magnetic resonance imaging. Nature 653, 411–417 (2026).

56. Tsukada, H., Nishiyama, S., Fukumoto, D., Kanazawa, M. & Harada, N. Novel PET probes 18F-BCPP-EF and 18F-BCPP-BF for mitochondrial complex I: a PET study in comparison with 18F-BMS-747158-02 in rat brain. J Nucl Med 55, 473–480 (2014).

57. Mansur, A., et al. Test-retest variability and reference region-based quantification of (18)F-BCPP-EF for imaging mitochondrial complex I in the human brain. J Cereb Blood Flow Metab 41, 771–779 (2021).

58. Das, S.R., Avants, B.B., Grossman, M. & Gee, J.C. Registration based cortical thickness measurement. Neuroimage 45, 867–879 (2009).

59. Avants, B.B., et al. The optimal template effect in hippocampus studies of diseased populations. Neuroimage 49, 2457–2466 (2010).

60. Foulon, C., et al. Advanced lesion symptom mapping analyses and implementation as BCBtoolkit. Gigascience 7, 1–17 (2018).

61. Ashburner, J. & Friston, K.J. Unified segmentation. Neuroimage 26, 839–851 (2005).

62. Glasser, M.F. & Van Essen, D.C. Mapping human cortical areas in vivo based on myelin content as revealed by T1- and T2-weighted MRI. J Neurosci 31, 11597–11616 (2011).

63. Sandrone, S., et al. Mapping myelin in white matter with T1-weighted/T2-weighted maps: discrepancy with histology and other myelin MRI measures. Brain Struct Funct 228, 525–535 (2023).

64. Glasser, M.F., et al. The minimal preprocessing pipelines for the Human Connectome Project. Neuroimage 80, 105–124 (2013).

65. Andersson, J.L., Skare, S. & Ashburner, J. How to correct susceptibility distortions in spin-echo echo-planar images: application to diffusion tensor imaging. Neuroimage 20, 870–888 (2003).

66. Smith, S.M., et al. Advances in functional and structural MR image analysis and implementation as FSL. NeuroImage 23, 208–219 (2004).

67. Dell’Acqua, F., et al. A model-based deconvolution approach to solve fiber crossing in diffusion-weighted MR imaging. IEEE transactions on bio-medical engineering 54, 462–472 (2007).

68. Guo, F., Leemans, A., Viergever, M.A., Dell’Acqua, F. & De Luca, A. Generalized Richardson-Lucy (GRL) for analyzing multi-shell diffusion MRI data. Neuroimage 218, 116948 (2020).

69. Daducci, A., et al. Accelerated Microstructure Imaging via Convex Optimization (AMICO) from diffusion MRI data. Neuroimage 105, 32–44 (2015).

70. Esteban, O., et al. fMRIPrep: a robust preprocessing pipeline for functional MRI. Nature Methods 16, 111–116 (2019).

71. Reprint of: Mahalanobis, P.C. (1936) “On the Generalised Distance in Statistics.”. Sankhya A 80, 1–7 (2018).

72. Yan, C.-G., Wang, X.-D., Zuo, X.-N. & Zang, Y.-F. DPABI: Data Processing & Analysis for (Resting-State) Brain Imaging. Neuroinformatics 14, 339–351 (2016).

73. Tononi, G., Edelman, G.M. & Sporns, O. Complexity and coherency: integrating information in the brain. Trends Cogn Sci 2, 474–484 (1998).

